# Systematic computational assessment of atrial function impairment due to fibrotic remodeling in electromechanical properties

**DOI:** 10.1101/2025.06.24.661244

**Authors:** Åshild Telle, Ahmad Kassar, Nadia Chamoun, Romanos Haykal, Alejandro Gonzalo, Tori Hensley, Yaacoub Chahine, Oscar Flores, Juan C. del Álamo, Nazem Akoum, Christoph M. Augustin, Patrick M. Boyle

## Abstract

Cardiac fibrosis is a pathological condition associated with many cardiovascular diseases. Atrial fibrosis leads to reduced atrial function, resulting in diminished blood flow and an increased risk of stroke. This reduced function arises from altered myocardial electrophysiological and mechanical properties. Identifying the relative importance of these fibrosis-associated properties can reveal the most significant determinants of left atrial function impairment.

In this study, we used a computational framework to investigate the relative importance of various fibrosis-associated properties. Our model, a 3D electromechanical framework coupled with a 0D circulatory model, incorporated patient-specific geometries and fibrosis distributions from clinical imaging data. Nine parameters related to fibrotic remodeling (conduction velocity, ion channel expression levels, cell- and tissue-scale contractility, and stiffness) were analyzed using two sensitivity analysis schemes: a one-factor-at-a-time setup, allowing for the analysis of isolated effects, and a fractional factorial design, enabling the examination of combined effects. As output, we tracked various metrics derived from model-predicted pressure-volume loops.

Impairment of L-type calcium current (I_CaL_) was most detrimental (up to 64% reduction in A-loop area). Conversely, reduced inward rectifier current (I_K1_) led to improved atrial function (up to 27% increase in A-loop area). Fractional factorial design analysis revealed that combination with other parameter changes blunted the impact of reduced I_CaL_ but amplified the impact of reduced I_K1_. Further analysis of spatiotemporal distributions linked these effects to changes in intracellular calcium handling.

Future research focusing on I_K1_ and I_CaL_ could be highly significant for clinical and scientific advances. Modeling work can potentially help evaluate left atrial function among larger patient cohorts, focusing on strain analysis. Our work could also be extended to spatiotemporal simulations of blood flow and thrombosis, shedding light onto the mechanisms underlying atriogenic stroke.

**Author summary:** Cardiac fibrosis is a process where healthy heart muscle is replaced with non-conductive, non-contractile tissue. This change disrupts how the heart beats and contracts. In the left atrium, fibrosis is strongly linked to atrial fibrillation and a higher risk of stroke, the latter due to impaired pumping and altered blood flow.

In this study, we used a detailed computer model of the heart, based on real patient-specific left atrial shapes and fibrosis patterns, to understand how different fibrosis-related changes affect atrial function. We tested nine features of the heart’s electrical and mechanical behavior that are known to change during fibrosis, aiming to identify which ones have the most impact on the atrial function.

We found that reducing the L-type calcium current — an important signal for muscle contraction — caused the greatest decrease in atrial performance. Surprisingly, reducing the inward rectifier potassium current actually improved it. These effects were tied to changes in calcium handling inside heart cells. Our findings highlight promising directions for future heart disease research and treatment.

## 1 Introduction

Cardiac fibrosis is prevalent in cardiovascular disease and contributes to left atrial (LA) dysfunction. LA fibrosis is strongly associated with atrial fibrillation (AF) and ischemic stroke [1–3]. Fibrotic remodeling encompasses a series of complex pathological events involving myocyte death, expansion of the extracellular matrix, and subcellular electromechanical changes [4, 5]. These alterations reduces LA function, which can be quantified to support mechanistic insight and clinical risk stratification.

Fibrotic remodeling profoundly impacts myocardial electrophysiological (EP) and mechanical properties. Structural tissue-level changes (myocyte necrosis) leads to decreased conduction velocity (CV) [6] and reduced myocardial force generation. Subcellular remodeling further reduces ion channel conductances [1, 5]. LA fibrosis is also linked to increased atrial stiffness [7], often attributed to up-regulated collagen crosslinking [8] and changes in collagen composition [9]; likely combined with myocyte stiffening [10–12]. These changes all affect the cardiac function, but our understanding of their relative contributions remains limited.

Computational modeling offers a powerful approach to elucidate the consequences of fibrosis-related alterations. By tuning parameters in physiologically informed models, one can predict consequences of specific pathological changes. Computational EP models of fibrotic LA have been used to study the connection between fibrosis and AF [13–16], revealing how altered CV and ion channel expression in fibrotic regions influence arrhythmia inducibility and spatial characteristics. Multi-physics, multi-scale modeling frameworks have been used to assess the impact of LA remodeling (including fibrosis) [17] and AF (without fibrosis) [18, 19] on pressure-volume (PV) relationships. Hemodynamic effects of fibrotic remodeling have also been explored via computational fluid dynamics analyses [20, 21]. All of these studies have shown utility in quantifying clinically relevant variables and enabling targeted investigations. Computational models offer an advantage over experimental and clinical studies in that the impact of different parameters can be isolated, helping to clarify their specific contributions and informing research priorities.

There are several approaches to make use of computational models to investigate individual impact of various parameters. One-factor-at-a-time (OFAT) analysis provides a systematic and straight-forward way to assess the isolated impact of each parameter [22, 23]; however, this does not account for combined or cooperative effects. Fractional factorial designs (FFD) setups offer a powerful method for reducing the parameter space by examining key combinations in a balanced and efficient manner [24, 25]. It allows for investigating individual and interactive parameter effects, while minimizing the number of required experiments. FFDs and design of experiment protocols have previously been used to study parameter sensitivity in ionic EP models [26, 27] and computational fluid dynamics analysis [28, 29]. Although FFD have been used only sparingly in cardiac computational modeling, the methodology hold potential for exploring large parameter spaces in detailed models of cardiac function [30]. To our knowledge, FFD has not yet been applied to multi-physics, organ-scale cardiac models.

This study combines computational modeling with OFAT and FFD analyses to disentangle the effects of various fibrosis-associated properties on atrial function. We performed simulations using three patient-specific LA geometries with corresponding fibrosis distributions, perturbing nine electromechanical parameters in fibrotic regions. We first conducted an OFAT sensitivity analysis to assess the isolated effect of each parameter. Next, we explored spatiotemporal distributions of membrane potential, intracellular calcium, and active tension from the simulations in which these parameters were changed to elucidate the mechanisms driving their importance. To gain deeper understanding of combined interactive effects, we performed a more detailed sensitivity analysis using a 2^9−5^ FFD scheme. To assess the effect of an increasing fibrosis burden, we repeated both analyses with 50% synthetically elevated fibrosis in the same three geometries. As output metrics, we tracked A-loop area, booster function, reservoir function, conduit function, and upstroke pressure difference during contraction, all derived from model-predicted PV loops. Sensitivity analyses based on these metrics were used to identify the most influential parameters.

## 2 Methods

### 2.1 Ethics statement

The study was approved by the Institutional Review Board of the University of Washington (STUDY00015081). A written statement of consent was obtained from each patient.

### 2.2 Patient recruitment

We obtained three patient-specific LA geometries with corresponding spatial distributions of fibrotic tissue, as well as electroanatomical mapping (EAM) data for the same patients. Participants were recruited from the University of Washington Medical Center, all of whom had AF and were scheduled for ablation. Exclusion criteria included prior atrial ablation, prior heart surgery, contraindications to late gadolinium enhancement MRI (LGE-MRI), pregnancy, gadolinium sensitivity, or inability to undergo MRI due to body mass or habitus constraints. Patient demographics, relevant LA clinical measurements, and derived patient-specific parameters are provided in Table 1.

**Table 1.** Patient-specific demographics, clinical measurements, and derived parameters.

|  | Patient 1 | Patient 2 | Patient 3 |
| --- | --- | --- | --- |
| Age | 81 | 63 | 70 |
| Sex | F | M | F |
| LA end-diastolic volume (mL) | 145.1 | 141.5 | 73.0 |
| LA fibrosis burden (%)* | 15.6 | 23.9 | 17.9 |
| Synthetically increased fibrosis burden (%)* | 23.4 | 35.85 | 26.85 |
| Total LA activation time (ms) | 82.0 | 56.0 | 116.8 |
| CV longitudinal direction ($CV_L$; cm/s) | 1.1500 | 1.5730 | 0.8100 |
| CV transverse direction ($CV_T$; cm/s) | 0.5142 | 0.7035 | 0.3614 |

\*of main LA body and appendage (excluding pulmonary veins)

### 2.3 Patient geometries and fibrosis distributions

LA geometries with corresponding fibrosis distributions were obtained from pre-ablation LGE-MRI images. MRI scans were taken at the end of atrial diastole (prior to contraction). Segmentation, processing, and analysis of raw MRI scans were performed by Merisight (Marrek Inc., Salt Lake City, UT), as previously described [31] and applied [15, 16, 32]. Fibrosis burdens are reported in Table 1 and annotated in Fig 1. The geometries were represented as 3D triangulated surfaces with fibrosis distributions (LGE maps), which we extruded by 2 mm [33, 34] outward to create volumetric geometries using CARPentry Studio (NumeriCor GmbH, Graz, Austria) [35].

**Fig 1.**
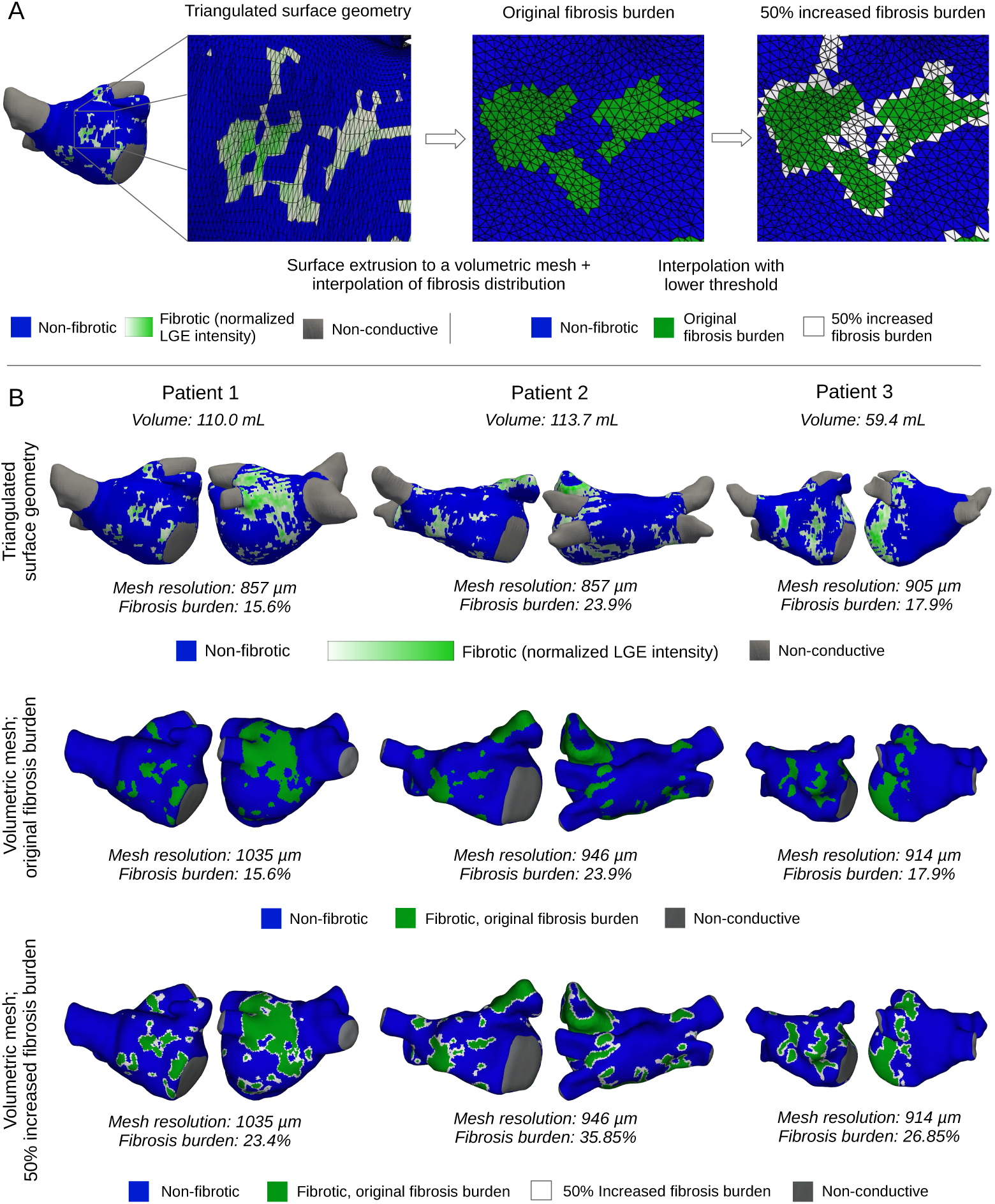
Patient-specific geometries with fibrosis distributions. (A) Workflow for mesh generation and fibrosis mapping: Starting with the original triangulated surfaces (left), we generated corresponding volumetric geometries. Fibrosis distribution was mapped via interpolation to preserve the original fibrosis burden (middle) and to reach a 50% synthetically elevated fibrosis burden (right). (B) Original shell geometries (top row), volumetric geometries with original fibrosis distributions (middle row), and volumetric geometries with increased fibrosis distributions (bottom row) for the three different patient geometries considered in this study. Mesh resolutions (average tetrahedral edge length) and fibrotic burdens are included as annotations.

Fibrosis distributions in the volumetric geometries were interpolated from LGE maps, ensuring that the total fibrotic burdens matched those reported by Merisight. All volumetric mesh elements were sorted and given a priority value according to the corresponding interpolated normalized LGE value. Following this order, we assigned elements one by one as fibrotic until the volumetric fibrotic ratio matched the original fibrosis burden. Fibrotic tissue was assumed to be uniformly distributed from the epicardium to the endocardium, with no transmural variation. Fibrosis burden was calculated as a percentage of mesh elements excluding the pulmonary veins (i.e., only the LA body and appendage), for original LGE maps and our volumetric geometries alike. To explore the effects of extended fibrosis, we applied the same method but with a target fibrosis burden increased by 50% (×1.5). Fig 1A illustrates how fibrotic regions were mapped from LGE maps to volumetric geometries for both levels of fibrosis, while Fig 1B displays the resulting volumetric geometries, with corresponding fibrosis distributions, for all three patients.

Volumetric geometries were further augmented as needed for our electromechanical simulations. Pulmonary veins were assumed passive post-ablation and hence not included as LGE map distributions. However, they were included in our modeling geometries without corresponding fibrosis maps. Artificial pulmonary vein caps and a mitral valve representation were added to the geometry to define boundary conditions and preserve anatomical orifice shape even under large deformations. Fiber direction maps (delineating the longitudinal myocyte directions) were generated for each geometry using a rule-based method, similar to those described in previous publications [35, 36].

### 2.4 CV calibration and identification of earliest activation locations

EAM was performed and recorded for all patients during the ablation procedure, yielding LA activation maps in sinus rhythm. A CARTO (J&J MedTech) system was used to obtain the values during the procedure, and the open-source OpenEP package [37] was used to extract local activation maps post-procedure. EAM data was analyzed to identify each patient’s earliest LA activation sites (first 5 ms) and total activation time.

Total activation times were subsequently used to personalize organ-scale CV values through an iterative inversion procedure. In this procedure, we fixed the anisotropy ratio for longitudinal CV (CV_L_) versus transverse CV (CV_T_) at 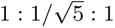 [14, 15], limiting the optimization to a single varying parameter (CV_L_). The optimization was performed by running EP simulations with our computational model, comparing the total activation time from the simulation to the recorded total activation time, and then adjusting the model’s CV values accordingly. This process was repeated until the difference in recorded and model-predicted activation times was less than 1 ms.

Activation times, selected electrical stimulus locations, and resulting simulated activation are displayed in Fig 2, while resulting CV values are listed in Table 1. Additional details are provided in S1 Appendix, including more detailed descriptions of the procedure (§ A.1) and extended figures showing activation and electrical stimulus locations from multiple views (Fig A1–A3).

**Fig 2.**
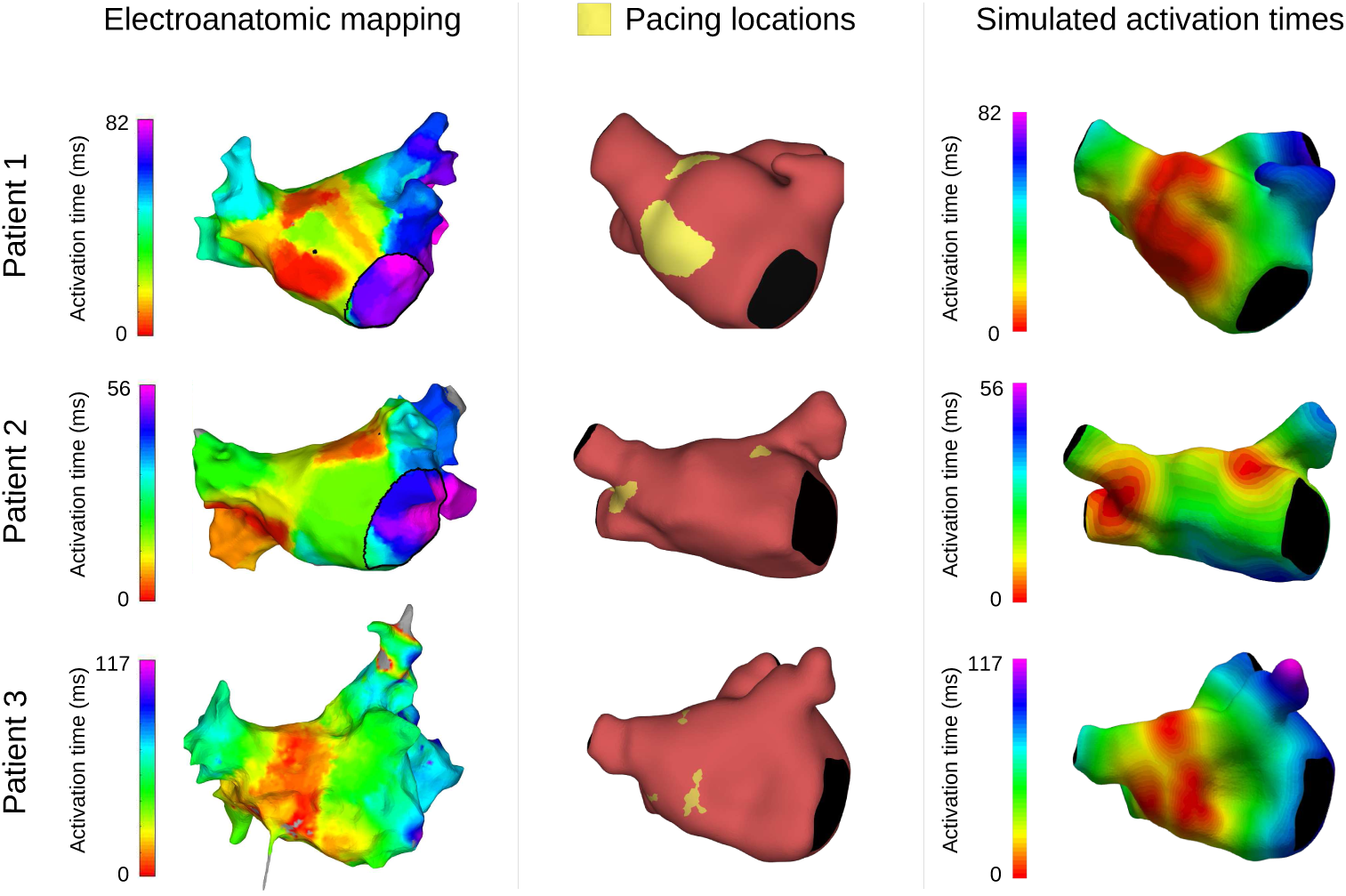
Electroanatomical mapping (EAM) activation time, electrical stimulation locations, and model activation times. Activation times estimated from EAM data (left), electrical stimulus locations corresponding to areas with the earliest activation times (first 5 ms; middle), and resulting activation times as simulated by our EP model (right). All activation times are displayed relative to first LA activation.

### 2.5 Multi-scale, multi-physics LA modeling framework

To predict changes in atrial function under different conditions, we employed a multi-scale, multi-physics physiological LA modeling framework. Electrical activation and mechanical contraction were modeled using a weakly coupled 3D electromechanical framework [21, 38, 39], with deformation strongly coupled to a 0D circulatory model [40]. Pulmonary veins were assumed to share the properties as non-fibrotic myocardium, exhibiting relative shortening within literature ranges [41].

Electrical activation was first modeled through a multiscale EP model, coupling cell- and tissue-scale dynamics. The cell-scale model was based on the human atrial action potential by Courtemanche et al. [42] with modifications according to Bayer et al. [43]. We simulated tissue-level electrical propagation using a reaction-eikonal model [44] with diffusion. Personalized CV values derived from EAM data (as described in § 2.4) were prescribed; baseline values are listed in Table 1. Pulmonary vein caps and the mitral valve were modeled as an in-excitable and non-conductive material.

The intracellular calcium distributions predicted by the EP framework was used as input to the biomechanical model. Active tension generation caused by intracellular calcium was computed using the cell-level contraction model developed by Land et al. [45, 46]. Maximum active tension (*T_a_*) was scaled with 50 kPa at baseline, calibrated to give approximately a 30% active emptying fraction [17, 47, 48]. Passive tension was modeled using a reduced Holzapfel-Ogden formulation with dispersion, with default material parameters *a* = 2.92 kPa, *b* = 5.6, *a_f_*= 11.84 kPa, *b_f_* = 17.95, and *δ_f_*= 0.09 [21, 39]. Pulmonary caps and the mitral valve were modeled as a passive, stiff Demiray material [49], with material parameters *a* = 10 000 kPa, *b* = 5.6. Both materials were modeled as nearly incompressible, with bulk modulus *κ* = 650 kPa [21].

To account for the impact of blood flow pressure and load from ventricular contraction, the 3D mechanical model of the LA was strongly coupled [40] with the 0D circulatory model CircAdapt [50, 51]. Coupling between the 3D and 0D models was achieved by simulating 0D blood flow through the pulmonary veins into the LA and through the mitral valve into the left ventricle, thereby modulating the pressure applied to the endocardial surface of the 3D LA model. Additionally, active tension generated in the 0D left ventricle was used to model atrioventricular plane displacement by applying a traction boundary condition on the mitral valve annulus of the 3D model. Additional mechanical constraints not captured in the 0D CircAdapt framework were imposed using Robin-type boundary conditions: spatially varying, normal spring boundary conditions were applied to the epicardial surface [52], while omnidirectional spring boundary conditions were imposed at the pulmonary vein inlets.

### 2.6 Simulation details

Simulations were performed using the software *Carpentry* (Numericor GmbH, Graz, Austria) [38], which is built on extensions of the open-source EP platform *openCARP* [53]. All simulations were carried out on Hyak, the University of Washington’s high-performance computing cluster (on nodes with a Intel(R) Xeon(R) Gold 6230 CPU processor), in parallel using 40 processes. The steps taken to perform a full simulation are described below.

We initially performed an unloading-reloading procedure to ensure a physiologically accurate diastolic pressure configuration. This ensured that the reinflated, pressurized geometry matched the initial mesh generated from image data. We used a backward displacement algorithm to find an unloaded reference configuration [54], then reloaded the reference geometry with a prescribed diastolic pressure of 10 mmHg [17, 39].

To ensure physiologically reasonable initial states, we next performed electromechanical simulations at the cellular level using *openCARP* ‘s single-cell tool *bench*. These simulations were run for 50 cardiac cycles with a fine temporal resolution of 0.025 ms. Leveraging the significantly lower computational cost of cell-level simulations compared to full organ-scale models, this approach allowed efficient approximation of steady-state conditions. The resulting cellular states were then used to initialize the subsequent full organ-scale simulations.

Organ-scale simulations were then performed for ten cardiac cycles with the full 3D framework. The ten cycles were simulated to reach a convergent state (convergence plots for baseline simulations displayed in Fig 3; in the 10th cycle there was less than 2% difference for both pressure and volume), balanced with a reasonable running time (16–20 hours per simulation). End-diastolic volume decreased in the converged solutions (Fig 3, middle row). The PV loops predicted by the final cycle simulated were used for our subsequent analysis. For each cycle, activation was initiated by applying an electrical stimulus in earliest activation regions, determined by EAM data (as described in § 2.4). The remaining myocardial tissue was then activated through the propagation of the electrical signal, leading to atrial contraction and subsequently atrial relaxation as given by our modeling pipeline. We used imposed a 1 Hz frequency to mimic sinus rhythm. Time steps were discretized at 0.025 ms for the EP model and 0.5 ms for the coupled biomechanical-0D circulatory model.

**Fig 3.**
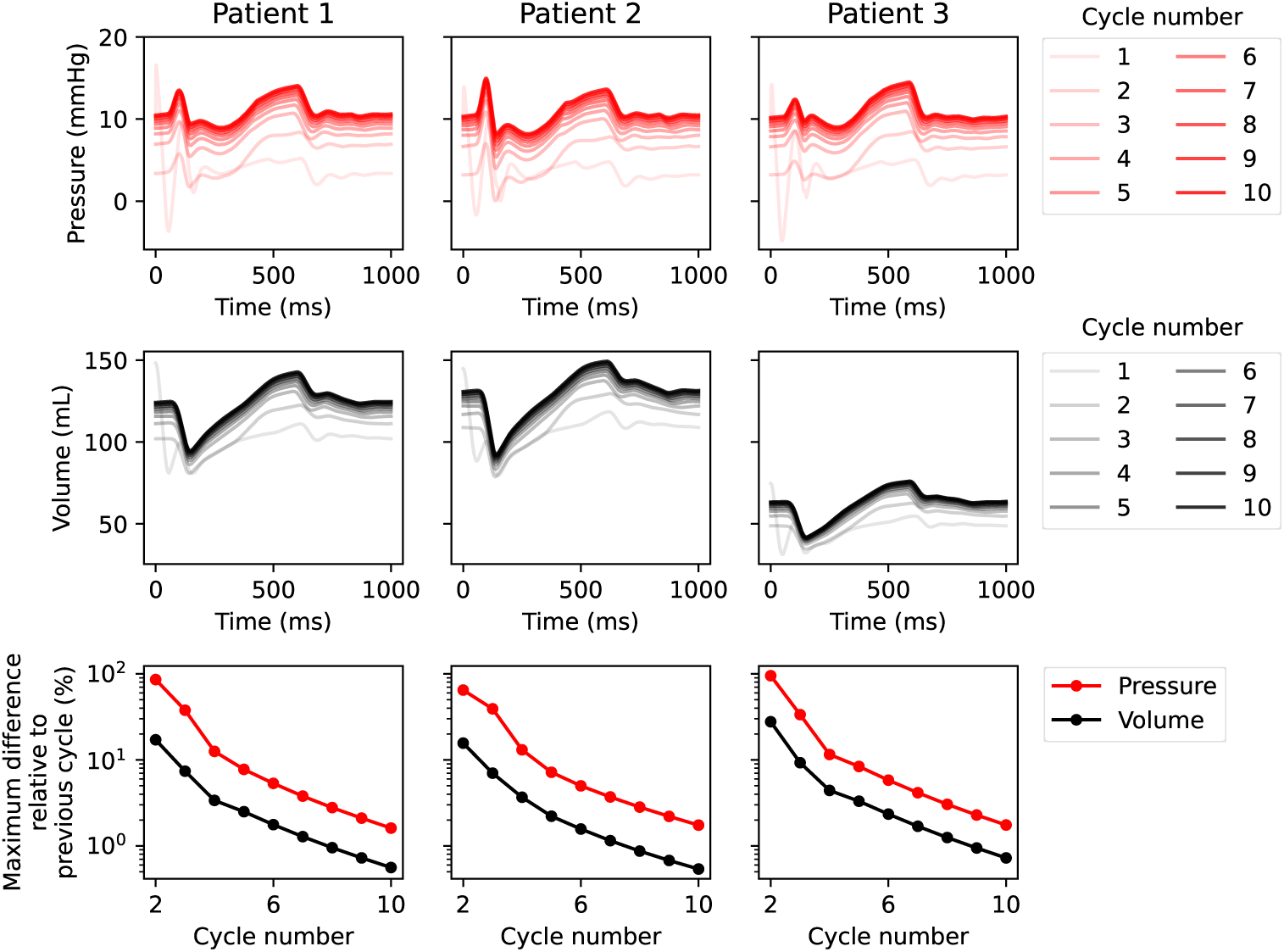
Convergence plots for baseline simulations. LA pressure (top) and volume (middle) over time over ten cardiac cycles; maximum differences in respective values between the current and previous cardiac cycles, normalized by the maximum value of the current cycle (bottom); for Patient 1–3, all at baseline.

### 2.7 Parameter changes in fibrotic regions

In our investigation of consequences of various fibrosis-related properties, we focused on five EP parameters (CV values and ion channel currents) and four mechanical parameters (contractile and passive properties); summarized in Table 2. Each parameter was re-scaled in the fibrotic regions by the corresponding factor (second column). Baseline or fibrotic parameter values were assigned in various combinations following the sensitivity analyses described in § 2.8. A complete overview of absolute values for all parameters in all combinations is included as Supplementary Data.

**Table 2.** Fibrotic remodeling parameter changes.

| Parameter | Scaling factor(s) | Notes | References |
| --- | --- | --- | --- |
| CV longitudinal direction ($CV_L$) | 0.657 | | [16] |
| CV transverse direction ($CV_T$) | 0.520 | Anisotropy ratio 1 : $1/\sqrt{8}$ | [14, 15] |
| Inward rectifier potassium current ($I_{K1}$) | 0.6 | Also alters CV values | [14, 15, 55–57] |
| L-type calcium current ($I_{CaL}$) | 0.5 | Also alters CV values | [14, 15, 55–57] |
| Fast sodium current ($I_{Na}$) | 0.5 | Also alters CV values | [14, 15, 55–57] |
| Myofilament binding rate ($\mu$) | 0.5 | | [58, 59] |
| Active tension scaling factor ($T_a$) | 0.5 | | [21, 60] |
| Increased longitudinal stiffness ($ST_L$) | $a \times 0.963$, $a_f \times 2.293$ | $2 \times$ load upon 5% stretch long. dir. | [7, 60] |
| Increased transverse stiffness ($ST_T$) | $a \times 2.157$, $a_f \times 0.675$ | $2 \times$ load upon 5% stretch transv. dir. | [7, 60] |
| ⇒Both; $ST_L$ and $ST_T$ in combination | $a \times 2.009$, $a_f \times 2.000$ | | |

For CV values, we independently explored reductions in the longitudinal and transverse directions. CV_L_ was scaled a factor of 0.657, based on values reported by Macheret et al. [16]. For CV_T_, we used a scaling factor of 0.520 (relative to baseline CV_T_ values); corresponding to a fibrotic anisotropy ratio of 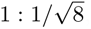, somewhat higher compared to the non-fibrotic ratio 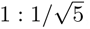; consistent with previous studies [14, 15].

Changes in ionic currents were imposed by scaling the inward rectifier potassium (I_K1_), L-type calcium (I_CaL_), and fast sodium (I_Na_) currents by factors of 0.6, 0.5, and 0.5, respectively [14, 15, 55–57]. These changes in ionic current levels are known to alter CV values locally [61, 62]. In a traditional tissue- and organ-scale EP model, these alterations would be captured indirectly through changes in membrane potential. However, the reaction-eikonal formulation we used in our study does not inherently account for the direct effects of ionic current changes; CV values are prescribed and not emergent properties of the underlying cell- and tissue-scale EP dynamics. To prescribe accurate CV values, we ran supplement simulations using OpenCarp’s *tuneCV* functionality [63]. These simulations were performed considering a simplified 1D rod to predict appropriate CV scaling factors for all combinations of ionic scaling factors. The resulting CV values for all possible parameter combinations, including in combination with reduced CV_L_ and CV_T_, are listed in(§ A.2, Table A1).

For contractile properties, we considered the active tension scaling factor (*T_a_*) and the myofilament binding rate (*µ*). Modifications to *T_a_* affect the magnitude of the active transient. We applied a fibrotic scaling factor of 50% for *T_a_* [21, 60] (relative to the baseline value of 50 kPa). The myofilament binding rate depends on the ratio of *α* and *β* myosin isoforms. This ratio shifts to a higher proportion of *β* myosins in AF [58, 59], indicating a slower contraction rate. We imposed a corresponding fibrosis-associated scaling factor of 50% (relative to the baseline value *µ* = 9 [46]) in our simulations.

To account for increased myocardial stiffness, we altered passive material parameters imposing changes in longitudinal (ST_L_) and transverse (ST_T_) stiffness. Specifically, we altered parameters *a*, which affects isotropic stiffness, and *a_f_*, which alters additional fiber direction stiffness. To consider stiffness independently for each direction, we performed virtual stretch experiments to estimate which values of *a* and *a_f_* were needed to achieve a two-fold change in the load (i.e., force per area) in one direction while keeping the load in the other direction unchanged. In these experiments, we stretched a tissue block (a unit cube) by 5% in either direction while tracking the corresponding load values, following the setup described in a previous publication [64]. Through an iterative procedure, we then stretched the tissue block, compared the load values with the original values, then altered the values of *a* and *a_f_*, and repeated the process until the load value doubled upon stretching in one direction while remaining unchanged in the other direction (within 1% precision). For the combined effect, having both ST_L_ and ST_T_ set to fibrotic levels, we found values of *a* and *a_f_* such that load values doubled when stretched 5% in either direction. The resulting *a* and *a_f_* values are listed in Table 2. Unloading-reloading procedure performed prior to simulations were done for each geometry at both levels of fibrosis and for all four possible stiffness configurations in fibrotic regions (without any changes; with increased ST_L_, with increased ST_T_, and both combined).

### 2.8 Sensitivity analysis – experimental setup

We investigated the impact of the nine identified fibrosis-associated parameters using two approaches. First, we used an OFAT analysis to analyze each parameter’s isolated effect. Next, a FFD analysis was used to evaluate combined effects. For this, we employed a 2^9−5^ design [65] as displayed in Table 3, resulting in 32 distinct parameter combinations. In this setup, B denotes the baseline factor level, while F represents the fibrosis-associated factor level, incorporating the relevant scaling factors (listed in Table 2).

**Table 3.**
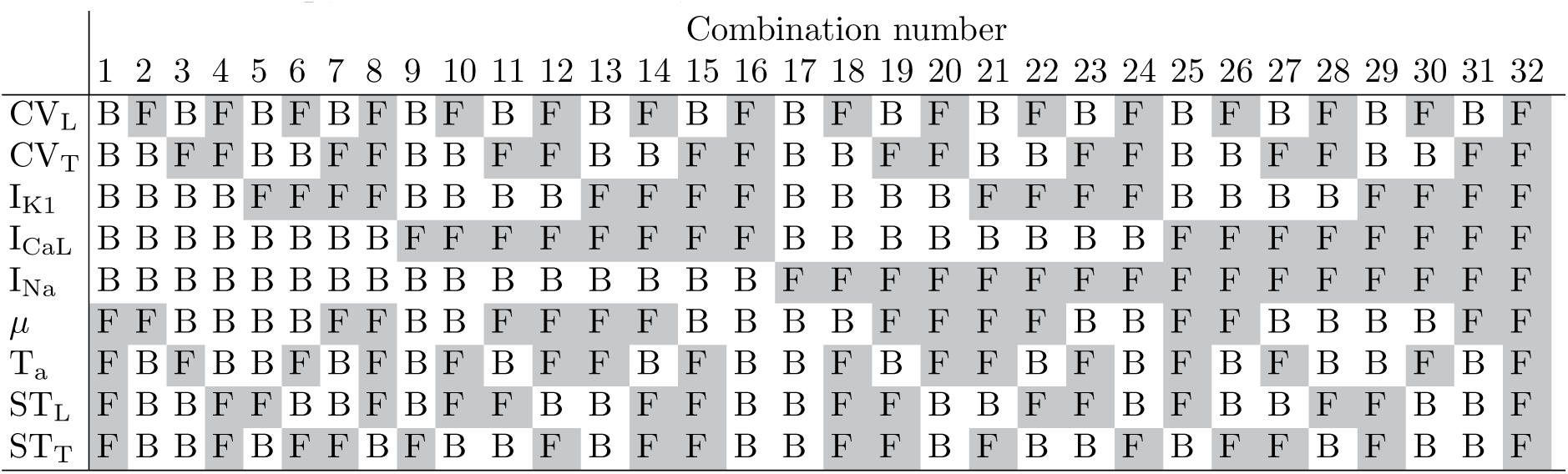
FFD setup; B = baseline value; F = fibrotic value.

### 2.9 Metrics reported

For all simulations, we report five PV-loop-based metrics capturing different aspects of atrial function. These include A-loop area [66–68], booster function, reservoir function, conduit function [69, 70], and upstroke pressure difference during contraction. The metrics are defined in the equations given below. Here, *LAV_preA_* and *LAP_preA_* refer to the LA volume and pressure at the the initial instant of electrical stimulus (LA end-diastolic volume), *LAV_min_* and *LAV_max_* refers to the minimal (LA end-systole) and maximal volumes, and *LAP_maxA_* refers to the maximal pressure during the A-loop (i.e., maximum value of booster phase pressure). We performed statistical analysis on values normalized by the corresponding baseline values for the same patient.

The five metrics considered were:

1. A-loop area (work performed during the active contraction): Calculated by Gauss’s area formula:

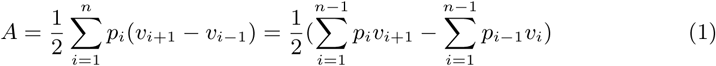 where *p* and *v* correspond to A-loop volume and pressure values.
2. Booster function (LA pumping function; active contraction volume change):

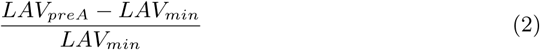
3. Reservoir function (elastic ability to stretch and recoil; passive stretching):

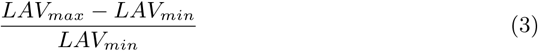
4. Conduit function (passive transfer of blood to the LV):

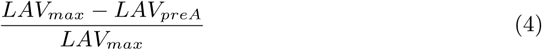
5. Upstroke pressure difference (LA pumping function; active contraction pressure change):

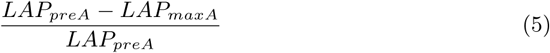

### 2.10 Spatiotemporal analysis

We examined spatiotemporal distributions of electromechanical simulations to better understand the effects of the parameters identified as most influential in our sensitivity analysis. Specifically, we compared the baseline simulation, those in which I_CaL_ and I_K1_ were impaired (from the OFAT analysis), and FFD Combination 32, in which all parameters were set to fibrotic levels. For conciseness, we refer to this combination as “Fully fibrotic” throughout the paper. We compared spatiotemporal distributions of membrane voltage, intracellular calcium, active tension, and fiber direction strain. This analysis was limited to the original fibrosis burden, with Patient 1 used as a representative example.

### 2.11 FFD analysis

To analyze the FFD main effect (isolated impact of a single parameter), we compared the baseline (B) and fibrotic (F) groups. For a given parameter, the B group included all combinations in which that parameter was set to baseline value, while the F group includes combinations where it was set to the fibrotic value (as defined in Table 2). Other parameters were set to either baseline or fibrotic levels according to the design in Table 3. Statistical comparisons between the two groups were performed using Student’s t-test (with group-wise distributions found to be normal, as assessed by an Anderson-Darling test); the analysis was conducted and annotated using the software Statannotations [71].

To characterize FFD interaction effects (combined or confounded effects with other parameters), the subdivision was extended to consider pairwise combinations. This resulted in four groups for each parameter pair (*x_i_, x_j_*) – all combinations of baseline/fibrotic levels (*B_i_B_j_*, *F_i_B_j_*, *B_i_F_j_*, and *F_i_F_j_*). We defined an interaction coefficient based on the *cross-product* between these:

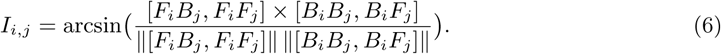

The coefficient relates the angle between two lines in an interaction plot. It is symmetric, meaning *I_i,j_* = *I_j,i_*. Positive values indicate a *synergistic effect*, where the impact of having both variables at fibrotic levels leads to a greater reduction in atrial function than the sum of the effects of individual factors. Negative values indicate *mitigating effect*, where the impact of having both variables at fibrotic levels leads to a smaller reduction in atrial function compared to the sum of the effects of individual factors.

## 3 Results

In this section, we present findings from both individual patient-specific simulations, using Patient 1 as a representative example, and statistical analysis of aggregated data. We begin by presenting an overview of the entire simulation sequence, PV loops with derived metrics for all three patients, followed by OFAT analysis results. We next present results from the spatiotemporal analysis focusing on parameter changes emerging as most important from the OFAT analysis, followed by FFD analysis results. Finally, we examine the impact of increased fibrosis. Unless otherwise noted, the results are representative of all three patients and reflect the original fibrosis burden.

### 3.1 Electromechanical simulations and PV loop-based metrics

Our modeling pipeline is demonstrated in Fig 4, for Patient 1 at baseline (no fibrotic changes applied). The model-predicted pressure and volume transients were extracted, resulting in characteristic LA PV loops. The PV loops corresponding to baseline simulations for all three patients are displayed in Fig 5, with derived metric values annotated. Patient 1 had the smallest booster and reservoir function. Patient 2 had the largest A-loop area and upstroke pressure difference, and the lowest conduit function (though only marginally compared to Patient 1). Patient 3 had the smallest A-loop area and upstroke pressure difference, but the highest booster, conduit, and reservoir function. No patient consistently showed higher metric values than the others.

**Fig 4.**
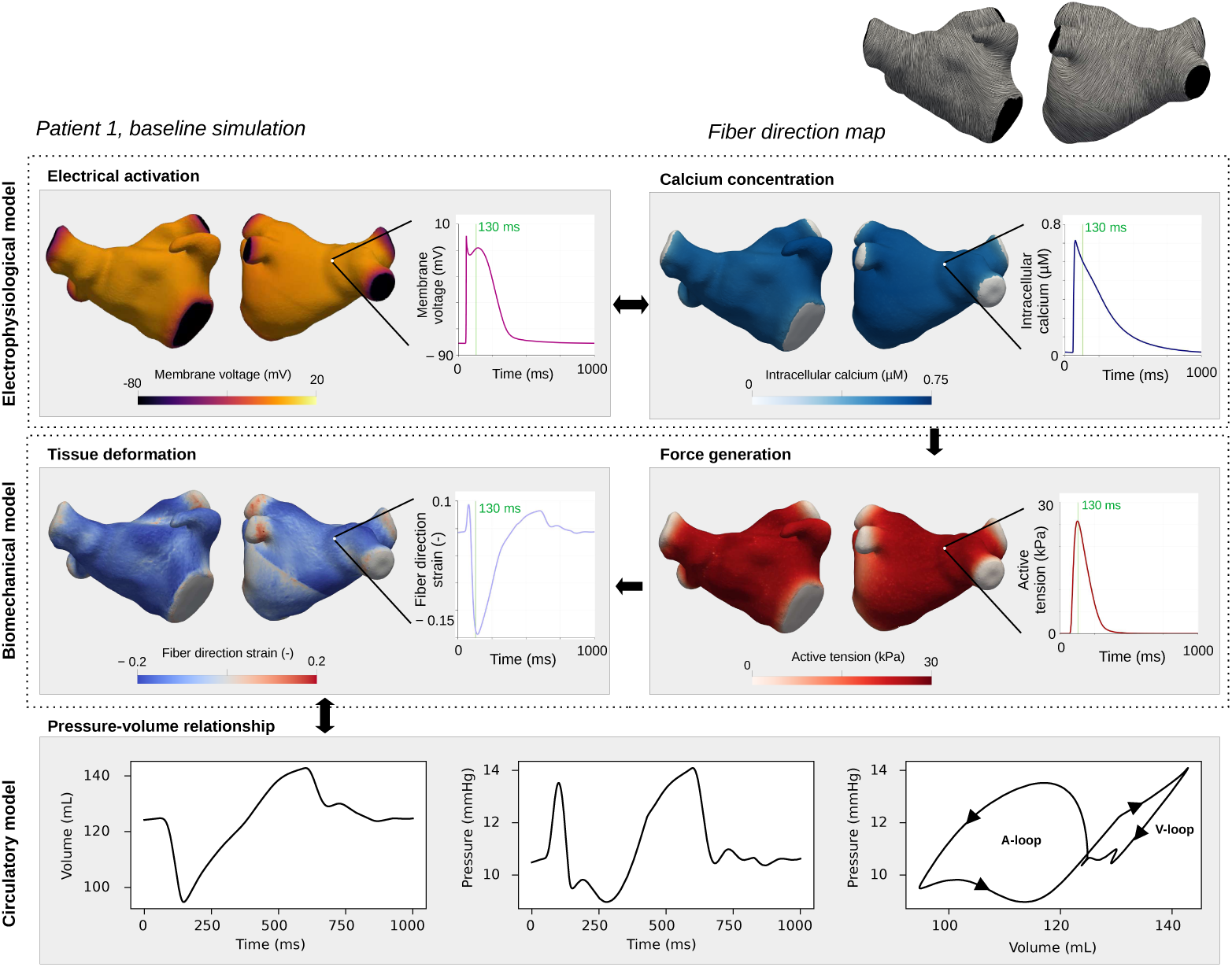
Model pipeline: EP, biomechanical, and circulatory models. Electrical propagation, including the release of intracellular calcium, was modeled with the EP model. The intracellular calcium resulted in the generation of active tension in our biomechanical model, which then caused tissue deformation. Tissue deformation was strongly coupled to the 0D circulatory model, from which we extracted LA volume and pressure transients. 3D maps (left) show the spatial distribution of model outputs at the 130 ms time point, and transient plots (right) display corresponding values over time at a representative node. Output data are shown for Patient 1, baseline simulation (no changes in fibrotic regions). Spatiotemporal distributions for all three patients are also displayed as movies included as supplementary material; see S2 Movie, S3 Movie, and S4 Movie.

**Fig 5.**
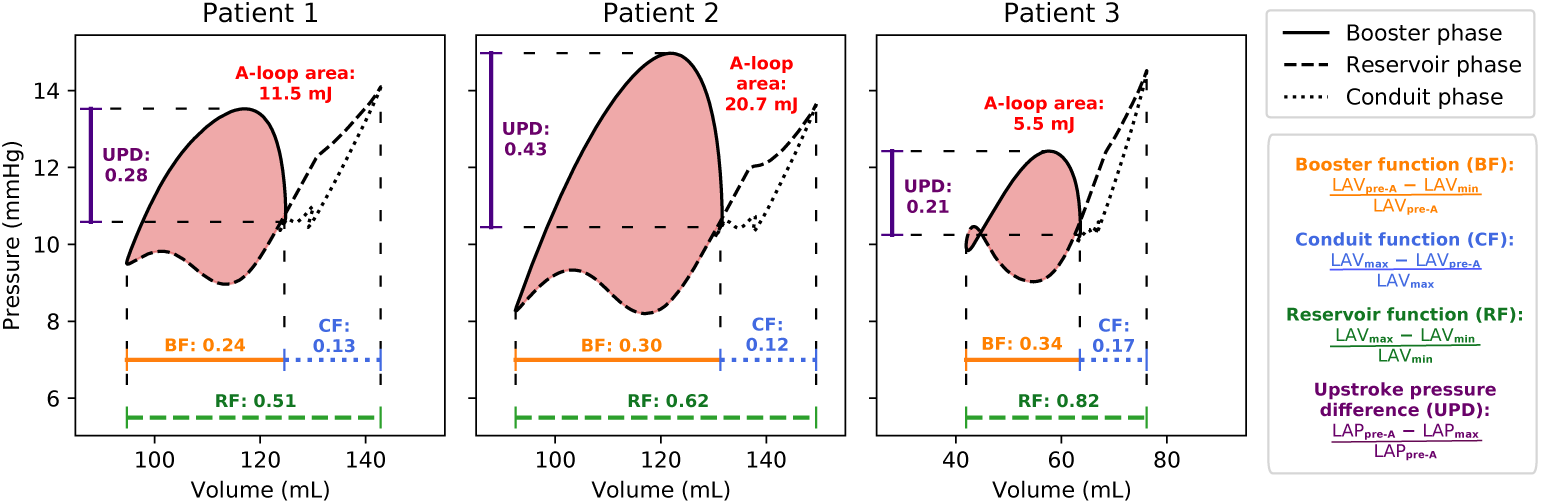
PV loops and derived metrics. PV loops from patient-specific simulations at baseline, for all three patients, with derived metrics annotated: A-loop area (i.e., stroke work in mJ), booster function (BF), conduit function (CF), reservoir function (RF), and upstroke pressure difference (UPD).

### 3.2 OFAT analysis predicted fibrosis-associated decreases in I_CaL_ and I_K1_ as main determinants of LA function

Results of our OFAT analysis are presented in Fig 6, highlighting the isolated effect of each parameter. Fig 6A shows the impact on PV loops using Patient 1 as a representative example. Impaired I_K1_ enlarged the A-loop, while impaired I_CaL_ (substantially) and reduced *T_a_* (marginally) decreased it. Increased stiffness (i.e., higher ST_F_ and ST_T_) shifted the PV loop, increasing volume while preserving pressure and loop shape. Modifying CV_L_, CV_T_, I_Na_, or *µ* did not noticeably alter PV loop size or shape.

**Fig 6.**
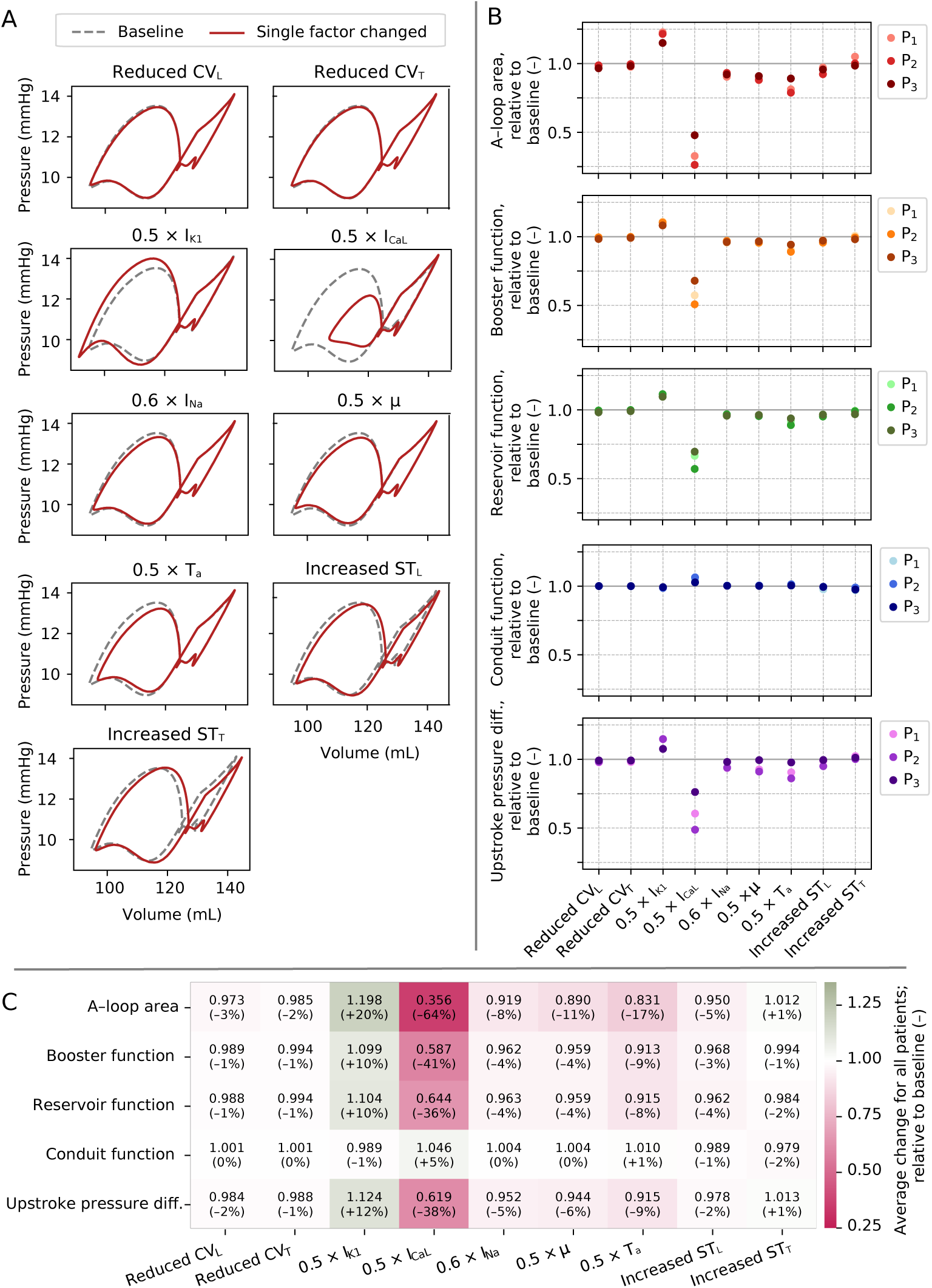
Results from OFAT analysis; original fibrosis burden. (A) PV loops with one parameter changed (in red) compared to baseline (gray, dotted), for Patient 1. (B) Impact of each parameter on A-loop area, booster function, conduit function, reservoir function, and upstroke pressure difference for Patients 1–3 (P_1_–P_3_). Values reported relative to baseline. (C) Average change (across all three patients) in each metric for each parameter, with implied percent-wise changes in parentheses.

Relative changes in PV-loop-derived metrics are plotted in Fig 6B. There were consistent trends across all three patients, with comparable magnitude changes. The largest changes were observed for Patient 2 (who had the highest fibrosis burden). A-loop area was the most sensitive metric (at maximum leading to a 74% reduction for impaired I_CaL_ for Patient 2).

Fig 6C displays output metrics and fibrosis-associated input parameters averaged across all three patients. Reducing I_CaL_ substantially decreased atrial function (with the largest decrease in A-loop area, 64%), while reducing I_K1_ increased function (with the largest increase in A-loop area, 20%). Reducing *T_a_* also had an effect (17% decrease in A-loop area). The impact of the other factors was otherwise modest; notably, reduced CV and increased stiffness did not alter any metric by more than 5%.

### 3.3 Impact of impaired I_CaL_ and I_K1_ was related to changes in intracellular calcium transient amplitude

Motivated by the results from the OFAT analysis, we next compared key spatiotemporal distributions across baseline, impaired I_CaL_, impaired I_K1_, and fully fibrotic simulations. Spatiotemporal distributions for all three patients under these conditions are also displayed as movies included as supplementary material; see S2 Movie, S3 Movie, and S4 Movie.

Fig 7 shows membrane potential, intracellular calcium, and active tension for each condition, using Patient 1 as a representative example. In the simulation with impaired I_CaL_, membrane potential was lower in fibrotic areas, while intracellular calcium was intermediate, and active tension was close to zero. A dispersive effect was also observed for calcium and active tension, where active tension in non-fibrotic areas was lower than in the baseline simulation (compare, e.g., the lower left region). In contrast, impaired I_K1_ abolished the differences between fibrotic and non-fibrotic regions producing a slight increase in intracellular calcium and a marked rise in active tension relative to baseline (see also transients in Fig 8). The fully fibrotic simulation exhibited pronounced heterogeneity in the intracellular calcium distribution, with near-zero values centrally and intermediate closer to non-fibrotic areas. Active tension was close to zero in all fibrotic regions, but higher in non-fibrotic areas compared to the I_CaL_ simulation.

**Fig 7.**
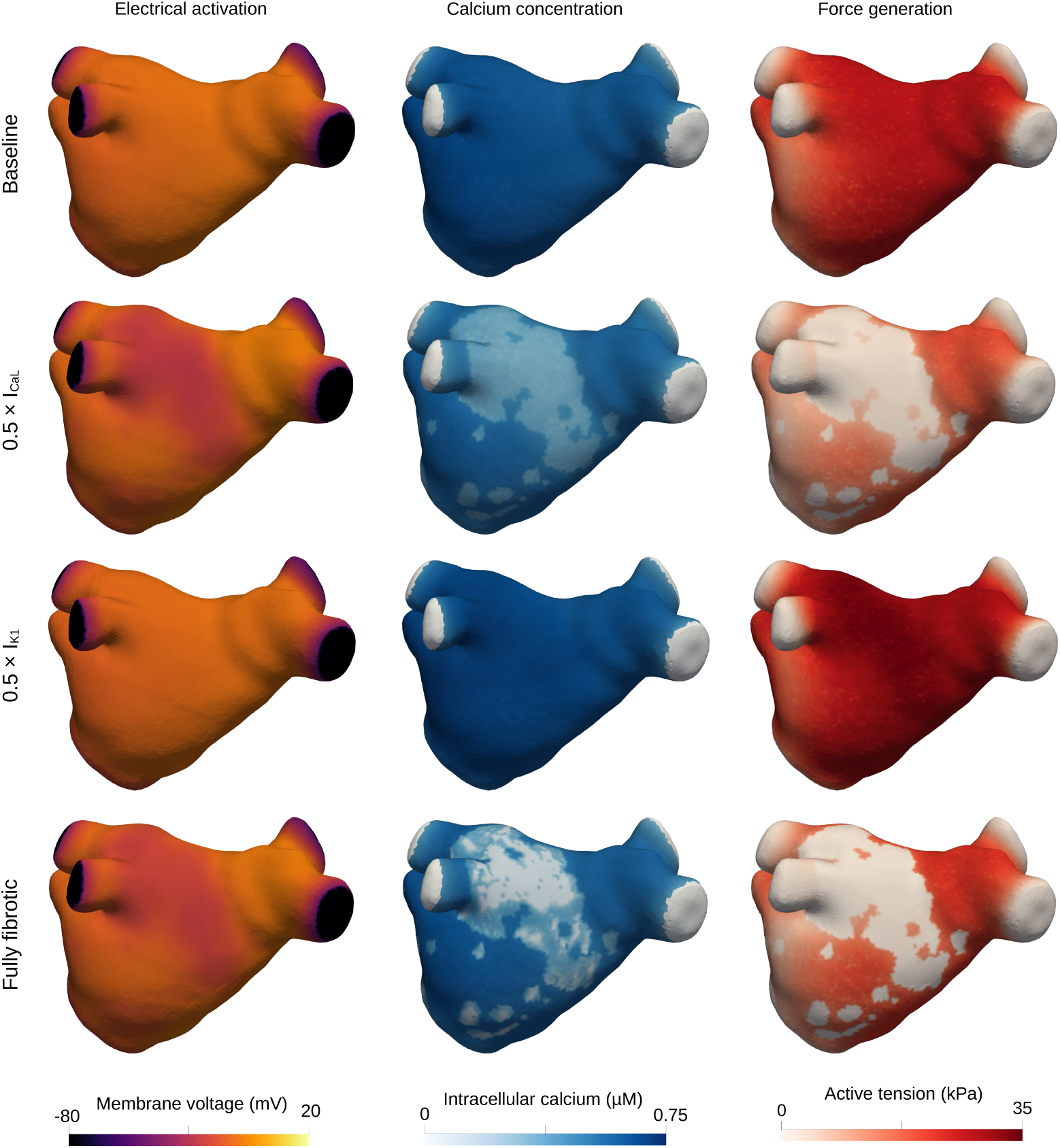
Spatial distributions of membrane potential, intracellular calcium concentration, and generated active tension; baseline, impaired I_CaL_, impaired I_K1_, and fully fibrotic simulations. Spatial distributions of membrane potential (left), intracellular calcium (middle), and active tension (right) at time step 130 ms for the baseline (top), impaired I_CaL_ (second row), impaired I_K1_ (third row), and fully fibrotic (bottom row) simulations for Patient 1. See also corresponding point-wise plots displayed in Fig 8, as well as movies attached as supplementary material; S2 Movie, S3 Movie, and S4 Movie (for all three patients).

**Fig 8.**
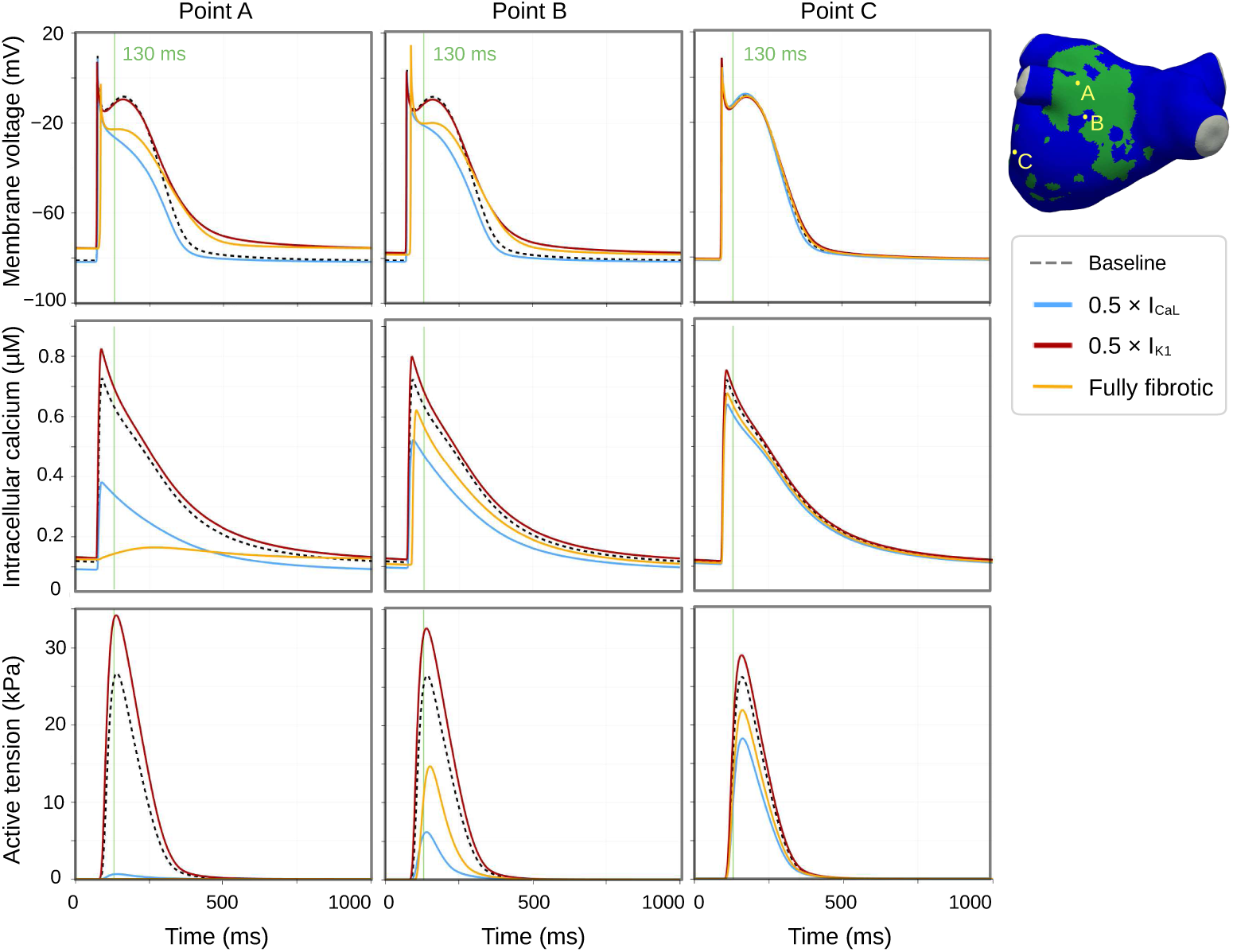
Membrane potential, intracellular calcium concentration, and generated active tension in selected points; baseline, impaired I_CaL_, impaired I_K1_, and fully fibrotic simulations. Membrane voltage, intracellular calcium, and active tension transients for three selected representative points (Point A – inside the fibrotic area; Point B – close to the fibrotic area; and Point C – away from the fibrotic area), Patient 1. Note that active tension for the fully fibrotic region is zero everywhere (not visible in the plot). See also corresponding spatial plots displayed in Fig 8.

Fig 8 shows membrane potential, intracellular calcium, and active tension transients for three representative locations in Patient 1’s geometry. The fibrotic site and its immediate vicinity (Points A and B) exhibited an elevated resting membrane potential (approximately 5 mV increase) and a prolonged action potential duration (approximately 75 ms longer) in both the impaired I_K1_ and fully fibrotic simulations. No substantial changes were observed at the distant site (Point C) for the membrane potential. The intracellular calcium transient amplitude was reduced for the impaired I_CaL_ simulation and slightly increased for the impaired I_K1_ simulation across all points; with gradual effect from Point A to Point B to Point C. In the fully fibrotic simulation, intracellular calcium was close to zero in Point A in the fully fibrotic simulation (see also corresponding spatial plots in Fig 7; bottom middle) while being somewhat reduced in Point B (14% decrease in maximum amplitude, relative to baseline) and Point C (7% decrease). These calcium variations magnified in active tension, with attenuated responses at lower calcium levels and enhanced responses at higher calcium levels. For the fully fibrotic simulation, active tension was zero in Point A, while decreased compared in Point B (44% decrease, compared to baseline) and Point C (19% decrease).

In Fig 9A, we show LA deformation. Deformation is displayed at the time of minimum volume (i.e., LA systole); see Fig 9B, top subplot). In the impaired I_CaL_ simulation (top row), the LA remained more dilated, indicating reduced contraction. Conversely, in the context of impaired I_K1_ (middle row), the deformed geometry was slightly more contracted than the baseline simulation. The fully fibrotic simulation (bottom row) also resulted in a less contracted geometry than in the baseline simulation.

**Fig 9.**
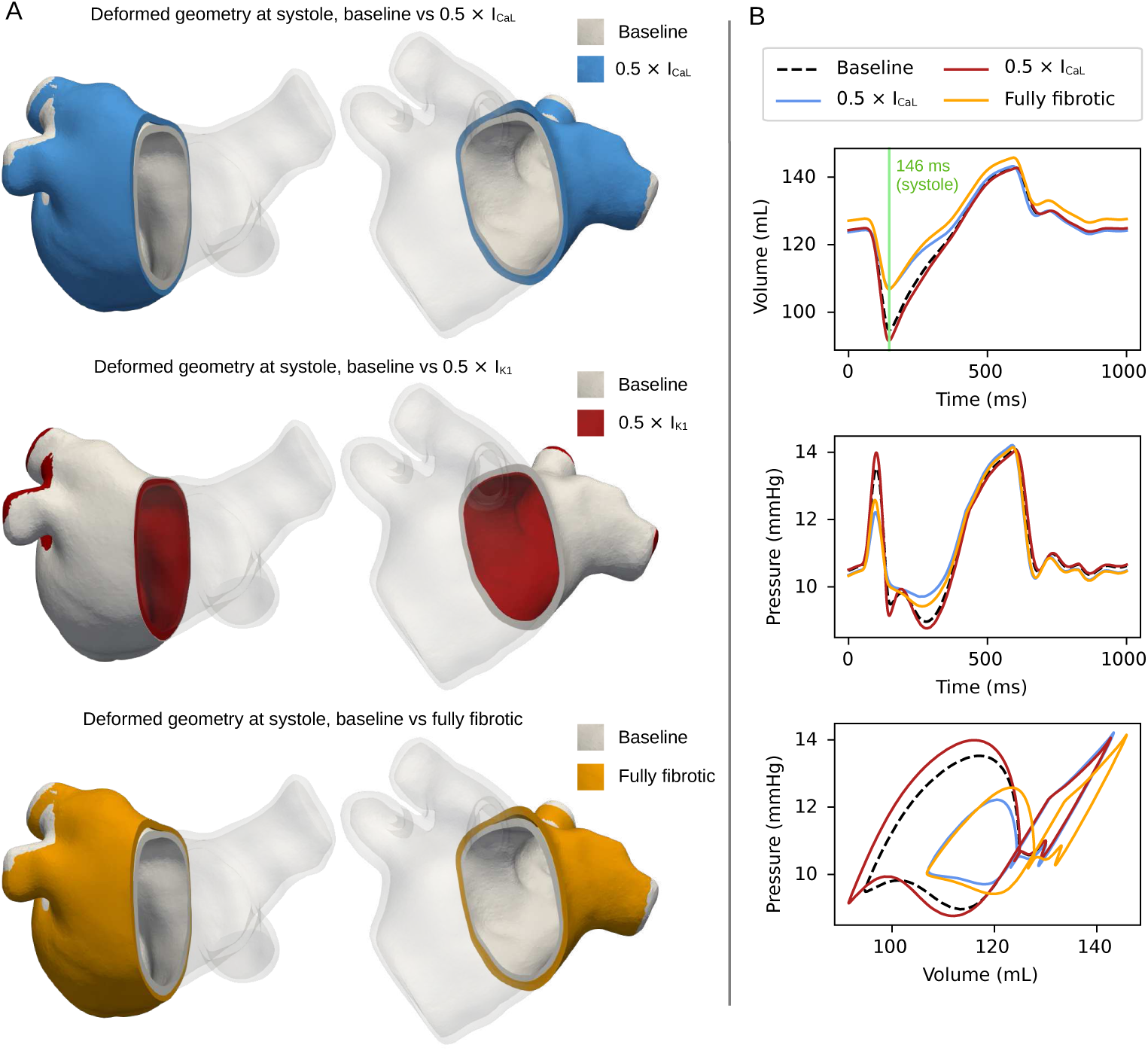
Geometry deformation and changes in volume and pressure; comparison of baseline with impaired I_CaL_, impaired I_K1_, and fully fibrotic simulations. (A) Deformed geometries for impaired I_CaL_ vs. baseline (top), impaired I_K1_ vs. baseline (middle), and fully fibrotic vs. baseline (bottom) simulations, Patient 1. Deformation is displayed from two different angles, with transparency applied to half of the geometry. (B) Volume over time, pressure over time, and PV loops for baseline, impaired I_CaL_, impaired I_K1_, and fully fibrotic simulations.

Fig 9B displays corresponding differences in volume and pressure over time. The systolic volume was similar between the impaired I_CaL_ and fully fibrotic simulations, while the fully fibrotic simulation exhibited a larger maximum volume. Pressure was also mostly changed during contraction, with the impaired I_CaL_ and fully fibrotic simulations having less variation than the baseline and impaired I_K1_ ones. The PV loop in the fully fibrotic simulations was correspondingly shifted in volume, and the A-loop was larger than in the I_CaL_ simulation.

### 3.4 FFD analysis predicted I_CaL_, I_K1_, longitudinal, and transverse stiffness as significant factors

To gain deeper understanding of the interactions between model parameters, we next performed a more detailed FFD analysis. The PV loops obtained from the simulations for all FFD combinations are shown in Fig 10, in which most combinations reduced the A-loop. Patient-averaged metric values for all combinations are included in S1 Appendix (Fig A4). For Combination 32 (fully fibrotic), the resulting relative change in A-loop area was 0.474 (i.e., −53%), in booster function 0.673 (−33%), in reservoir function 0.695 (−30%), in conduit function 1.0 (no change), and in upstroke pressure difference 0.721 (−28%).

**Fig 10.**
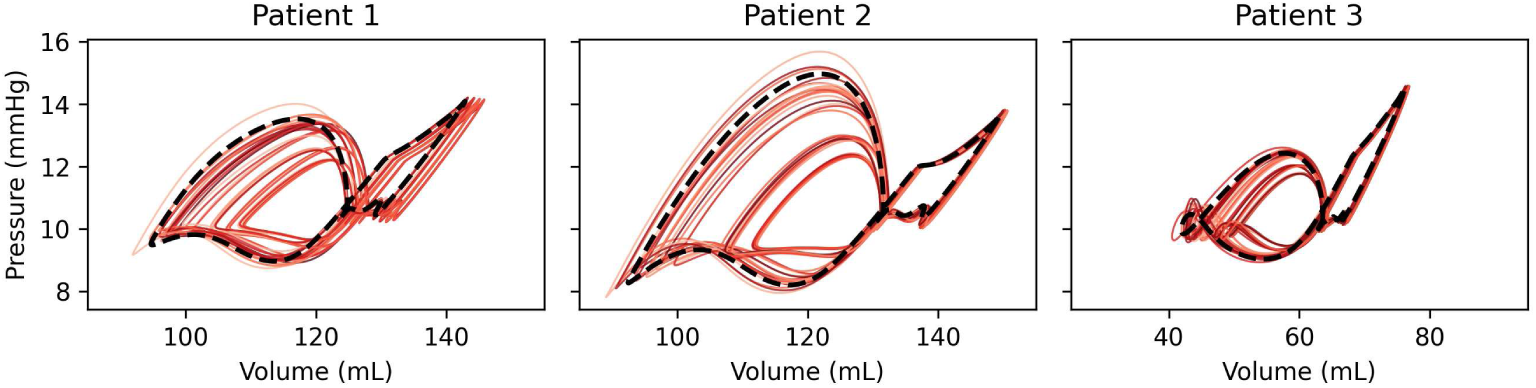
PV loops for all FFD combinations; original fibrosis burden. PV loops resulting from all FFD combinations, for all three patients. Baseline PV loops are displayed with a bolder black dotted line, while the thinner red traces correspond to various FFD combinations, as listed in Table 2.

Main effect plots for all patients and combinations are displayed in Fig 11A, with the relative change of each factor shown in Fig 11B. Reduction of I_CaL_ and I_K1_ both had a statistically significant impact on four of five metrics when comparing B and F groups; the impact in line with the OFAT analysis. The impact of impaired I_CaL_ was slightly attenuated (e.g., 54% reduction of A-loop area vs. 64% reduction for the OFAT analysis), whereas the impact of I_K1_ impairment was amplified (e.g., 27% increase of A-loop area vs. 20% increase for the OFAT analysis). Consistent with our OFAT analysis, changes in CV and stiffness values had a modest impact; here all less than 2%.

**Fig 11.**
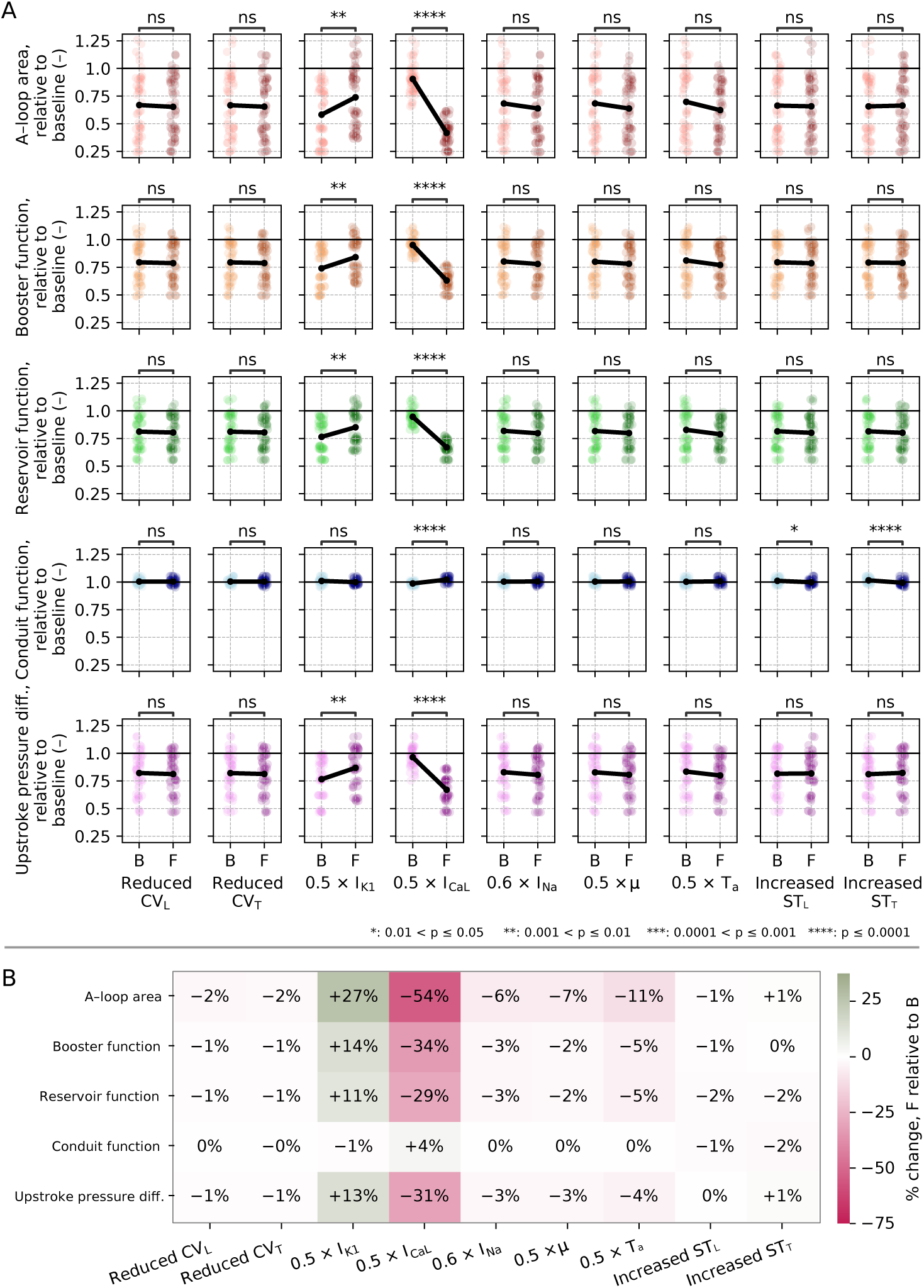
FFD – main effect; original fibrosis burden. (A) Main effect plots for each parameter and each metric, across all simulation combinations (Table 3) and all three patients combined. In each subplot, the combinations are divided into groups B or F based on whether the given parameter is set to baseline or fibrotic value. Points display values from individual simulations (i.e., all FFD combinations for all three patients), with lighter colors for the B group and darker colors for the F group. Lines show the average change from the B group to the F group. (B) Average increase/decrease across all patients and combinations for each metric listed numerically; calculated based on the difference between F and B, relative to B levels.

Increasing stiffness had a statistically significant albeit modest impact on conduit function, with decreases of 1% and 2% for increases in longitudinal and transverse stiffness, respectively. The impact was small in magnitude relative to the effects observed for the other metrics. These relative changes were consistent with little variation (all standard deviations fell within 0.03, and all observations were within the range [0.95, 1.06]); as such, statistically significant despite having a relative small impact.

Interaction effects presented in Fig 12 were generally small. All interaction coefficients (Fig 12A) were less than 0.1 radians, and both positive and negative interaction lines remaining nearly parallel (Fig 12B–C). Most combinations were found to be mitigating (having negative interaction values; see again Fig 12A). For A-loop area, booster function, reservoir function, and upstroke pressure difference, the greatest mitigating effect consistently was for reduced CV_T_ combined with increased ST_T_. Notably, there was also a mitigating effect for I_CaL_ combined with I_K1_; strongest for A-loop area, followed by booster function.

**Fig 12.**
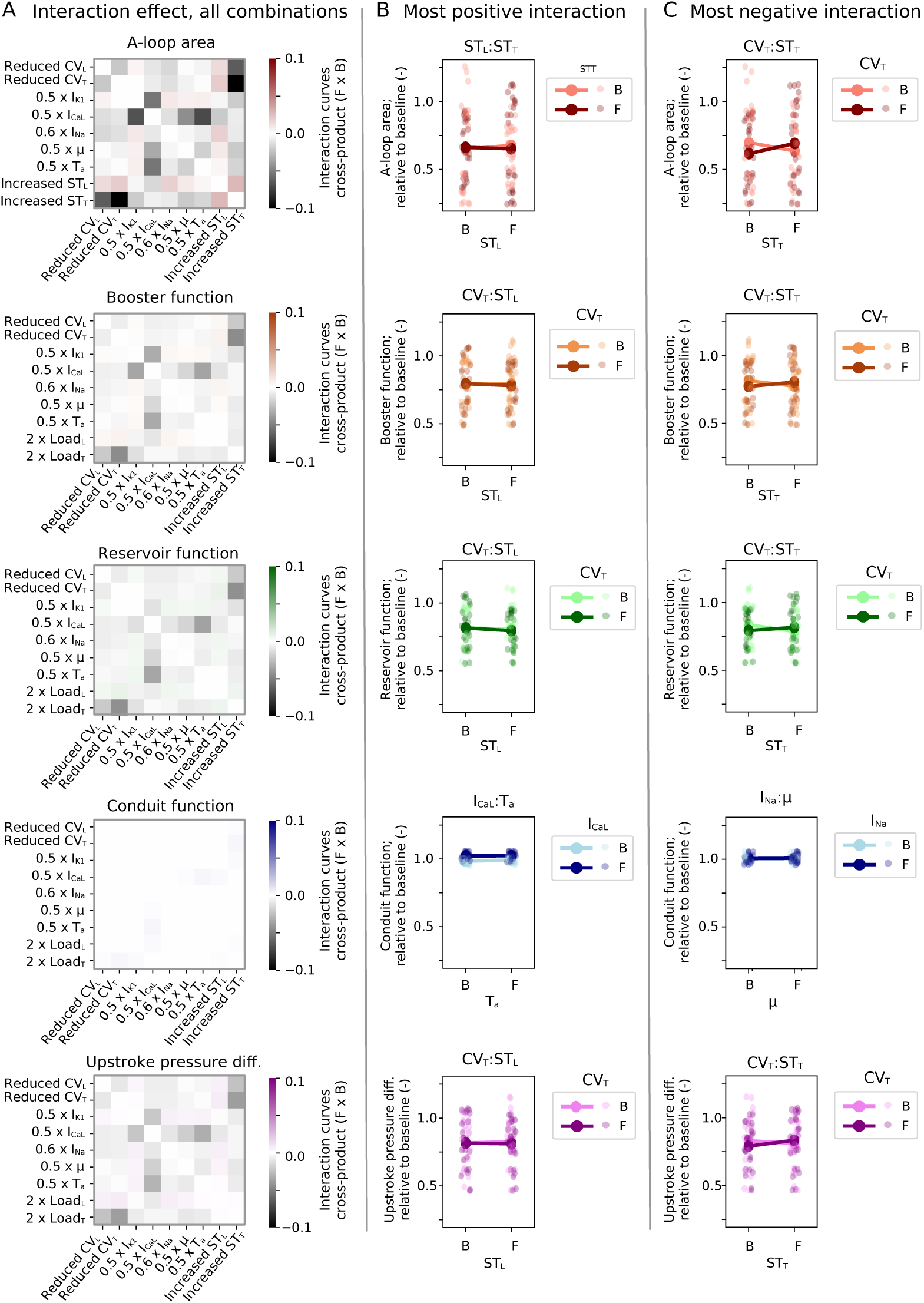
FFD – interactive effect; original fibrosis burden. (A) Interactive effects for each parameter pair, as calculated by Eq (6), across all simulation combinations (Table 3) and all three patients combined. (B–C) Interaction lines for the combination with the most positive (enhancing; B) and most negative (mitigating; C) impact. Points represent values from individual simulations (i.e., all FFD combinations for all three patients), while lines display average changes within subgroups.

### 3.5 Increased fibrosis led to a moderate further reduction in atrial function

Results of our analysis with a 50% synthetically elevated fibrosis burden are presented in Fig 13. Increased fibrosis burden led to a further reduction across all metrics in the OFAT analysis (Fig 13A), however, the decrease was moderate. Factors identified as statistically significant for FFD main effect subject to original fibrosis burden remained significant with elevated fibrosis (Fig 13B). Considering average values, (Fig 13C) the largest increase in absolute effect was observed for impaired I_CaL_. For A-loop area, we observed a 74% decrease with elevated fibrosis (compared to 64% with the original fibrosis burden) in the OFAT analysis, and a 62% decrease (compared to 54%) in FFD analysis.

**Fig 13.**
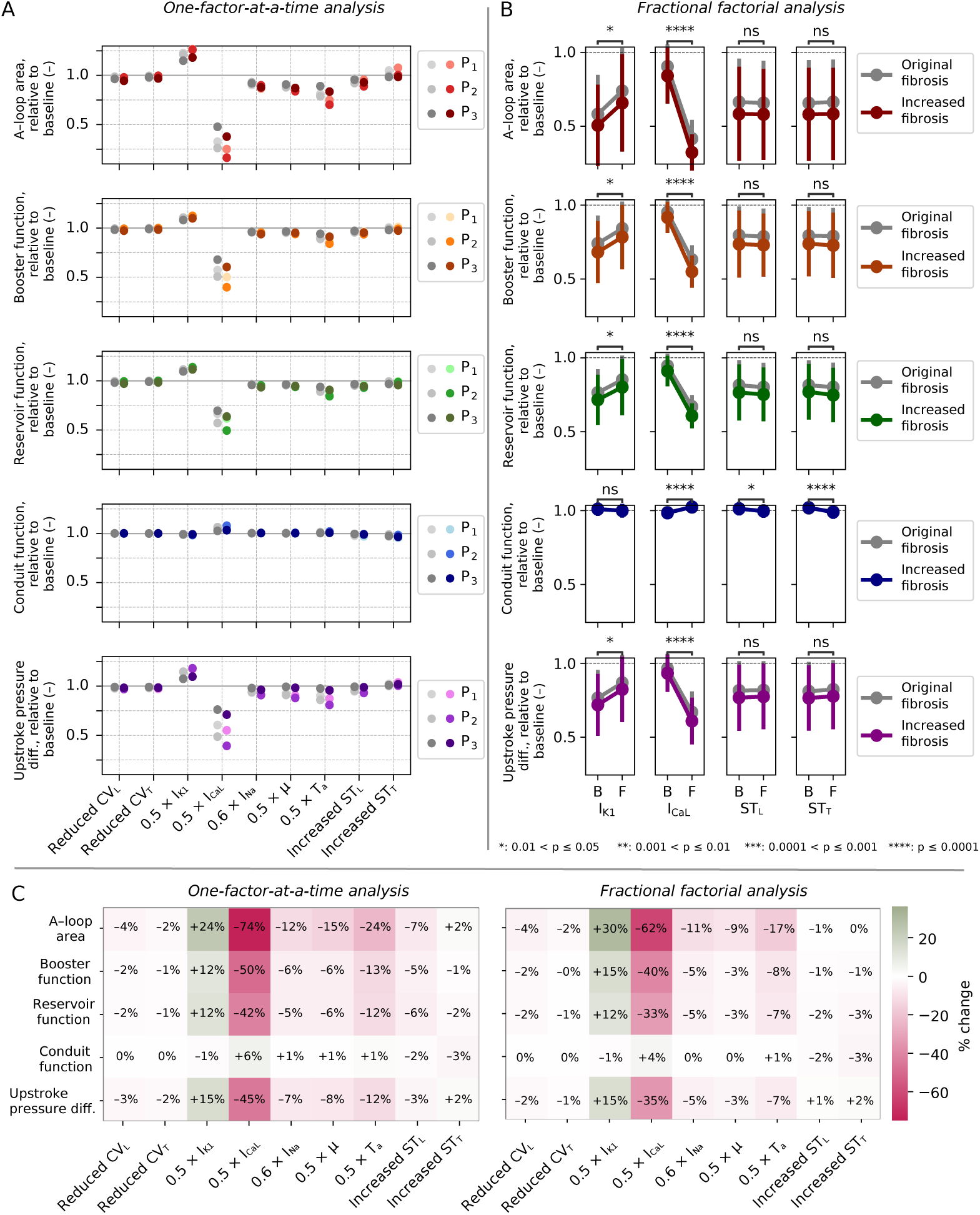
Impact of 50% synthetically elevated fibrosis; OFAT analysis and FFD analysis, main effect. (A) Results from the OFAT analysis for each patient; original fibrosis levels are indicated by gray points on the left (same as in Fig 10) and extended fibrosis on the right. (B) FFD main effect, including only columns with statistically significant differences. Error bars indicate standard deviation, and gray plots represent original fibrosis levels (same underlying data as in Fig 11). Comparisons for significant differences were performed for elevated fibrosis simulations. Only significant factors displayed; plots for all are included in S1 Appendix (Fig A6). (C) Percentage impact of each parameter, averaged across all three patient cases, for the OFAT analysis (left) and the FFD (right).

Patient-averaged metric values for all FFD combinations are included in S1 Appendix (Fig A5). For Combination 32 (fully fibrotic), the relative change in A-loop area was 0.366 (i.e., −63%), booster function 0.584 (−42%), reservoir function 0.622 (−38%), conduit function 0.995 (0.5% change), and upstroke pressure difference 0.657 (−28%).

In Fig 14 we display how absolute values of all metrics vary between no fibrosis, original fibrosis, and increased fibrosis burden; considering the fully fibrotic simulations. The decrease in all but the conduit function is steeper going from no fibrosis to original fibrosis burden compared to going from original to elevated fibrosis burden. Patient-specific trends correlated well between A-loop area and upstroke pressure difference, and between booster function and reservoir function, while conduit function differed from all other metrics.

**Fig 14.**
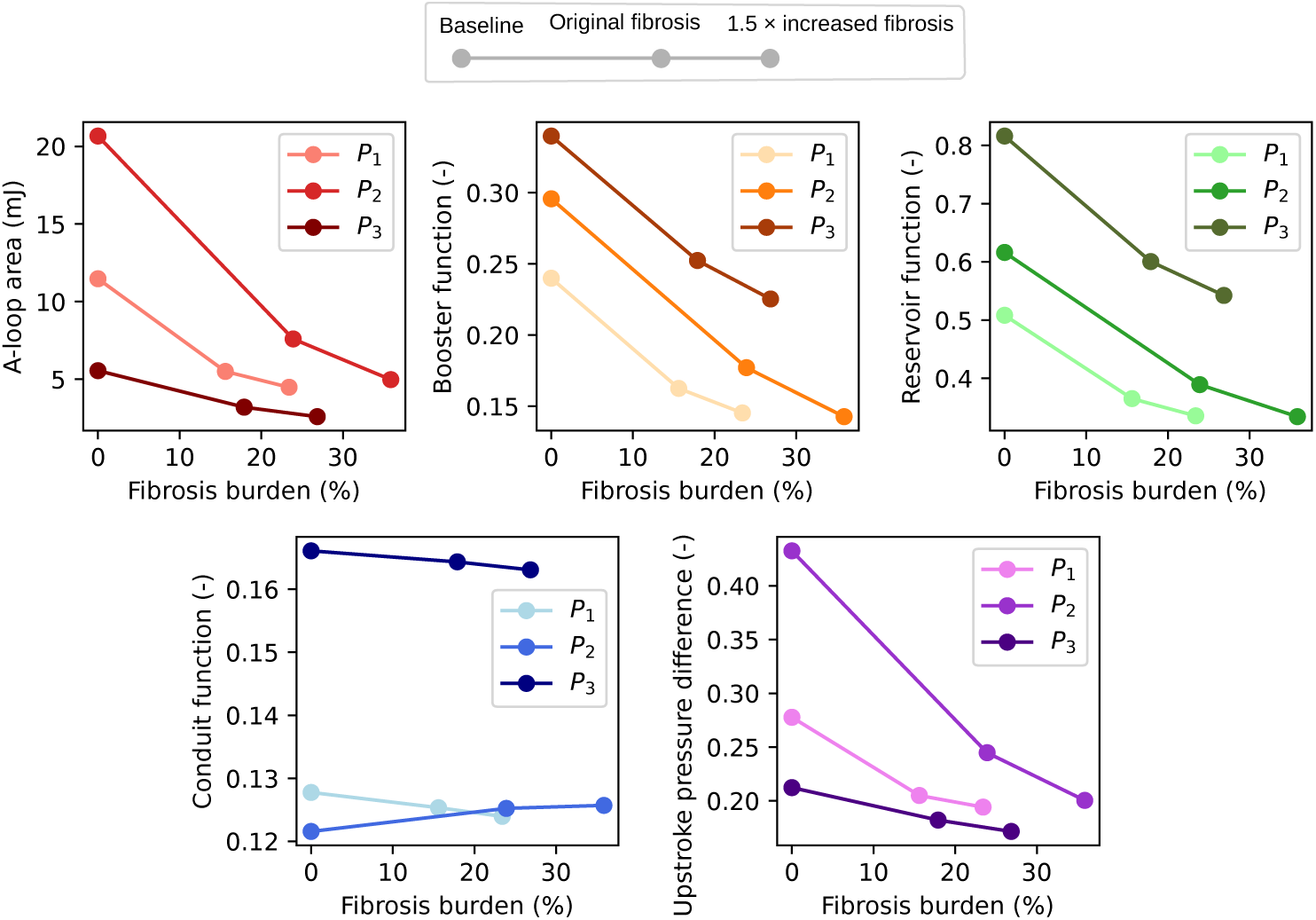
Absolute values of our five metrics for increasing levels of fibrosis; fully fibrotic simulation. Absolute values of A-loop area (stroke work), booster function, reservoir function, conduit function, and upstroke pressure difference for Patients 1–3 (P_1_–P_3_). For each patient, metrics are shown for baseline (0% fibrosis), original fibrosis burden, and 50% synthetically increased fibrosis burden.

## 4 Discussion

### 4.1 Impact of fibrotic-associated parameter changes on the atrial function

In the present study, we combined a multi-scale, multiphysics modeling framework with patient-specific LA geometries and corresponding fibrosis maps to investigate the theoretical impact of fibrotic remodeling. We investigated the influence of nine electromechanical parameters known to be altered by fibrosis, and assessed LA function through five metrics based on volume and pressure changes. We found that these metrics were sensitive to variations in I_CaL_ and I_K1_, with opposite effects, while stiffness exerted a small but statistically significant impact on conduit function. Spatiotemporal analysis revealed that subtle changes in intracellular calcium transient amplitude had pronounced effects on the active tension generated, reflecting the highly nonlinear relationship between these variables.

The impact of I_CaL_ on atrial function was consistent with its central role in regulating intracellular calcium release, which is essential for initiating cardiac contraction [72]. Atrial function remains partially impaired after sinus rhythm is restored in AF patients, primarily due to reduced L-type calcium current [73]. Less intuitive was the improved atrial function following impaired I_K1_. Experimental studies have demonstrated that increased intracellular calcium reduces the inward rectifying potassium current [74–76]; thus, we find it plausible that impaired I_K1_ could conversely raise intracellular calcium. To our knowledge, our study is the first delineating the impact of different fibrosis-associated changes on atrial function through electromechanical simulations on patient-specific geometries. Our results are, however, consistent with those of Hurtado et al. [77], who performed a sensitivity analysis on a simplified bar geometry comparing key parameters in two electromechanical ventricular models. They identified L-type calcium channel conductance followed by potassium delayed rectifier channel conductances as sensitive parameters. Increased L-type calcium channel conductance was found to increase intracellular calcium amplitude, while increased potassium delayed rectifier channel conductances conversely led to a decrease. This is completely in line with our results and previous experimental work.

In an extensive sensitivity analysis considering all four chambers of the heart, Strocchi et al. [78] found that transverse atrial stiffness (*b_t,A_*) had a more substantial impact than L-type calcium channel conductance on both end-systolic and end-diastolic LA volumes. On the other hand, they found potassium channel conductances to have little to no impact. The apparent discrepancy with our results might be explained by differences in sensitivity analysis parameterization and output metrics. Specifically, we observed that changes in stiffness shifted both end-systolic and end-diastolic LA volumes but not substantially their relative difference, i.e., the booster function. Next, the stiffness parameter in Strocchi et al. [78] is an exponent that with comparable fold changes have a much larger effect. The impact of stiffness has also been explored by others – Moyer et al. [17] found that fibrosis-associated increase in global stiffness reduced both the A-loop and the P-loop, while increasing pressure and reducing the volume in the passive phase. Meskin et al. [68] used silicone-based models, finding that compliance was correlated with lower variation in pressure and decreased A-loop area (stroke work). In our analysis, stiffness increased the LA volume also in the passive phase, while pressure and A-loop area remained largely unchanged. The positive shift in LA volume in our model likely arises from a change in mitral valve blood flow, in which higher stiffness led to a less efficient LA emptying. A shift in LA volume has also been observed in clinical studies [79], however, this is likely also due to LA remodeling. Despite differences in impact, previous studies and ours all support that stiffness impacts atrial function.

### 4.2 Sensitivity analysis takeaways

We performed two types of sensitivity analysis: a simpler OFAT analysis, which explored the isolated impact of each factor, and a more detailed FFD analysis, which also accounted for interactive effects. We found that most of the effects could be predicted by the OFAT analysis. However, FFD analysis revealed that the impact of I_CaL_ was partially mitigated when combined with other fibrotic-associated parameter changes, while the impact of I_K1_ was amplified. In the model, these changes can be linked to shifts in equilibrium values, observed through changes in action potential and calcium transient morphology (Fig 7). Physiologically, these might be considered compensatory feedback mechanisms.

Sensitivity analysis can can be performed in various ways, involving many methodological choices. Advanced techniques like the Morris elementary effects method [77, 80, 81] and Sobol indices [78, 82–84] involve finer sampling across predefined intervals, capturing non-linear behavior, interaction effects, and critical sensitivity regions. However, these benefits come at the cost of running more simulations, often requiring model simplifications or computational optimization strategies to manage feasibility. In our study, we sampled all relevant parameters at two levels: baseline and fibrotic. Nevertheless, the FFD scheme presented an efficient methodology allowing us to perform our sensitivity analysis taking into account interaction with a detailed multi-scale, multi-physics computational model based on a manageable number of simulations.

### 4.3 Clinical implications

We found that impairment of I_CaL_ in fibrotic regions had the most significant impact on atrial function, substantially reducing it, while reduced I_K1_ led to increased atrial function. With fibrotic area constituting between 15.6% and 23.9% (for original fibrosis burden), for the fully fibrotic simulation (with all factors set to fibrotic levels), A-loop area, booster function, reservoir function, and the difference in upstroke pressure decreased by 53%, 33%, 30%, and 28%, respectively. However, with 50% increased fibrosis burden, we only saw a very moderate further impairment. Our spatial analysis revealed substantial diffusive impact from the fibrotic to the non-fibrotic regions. The moderate effect could be explained by the fact that synthetically elevated fibrosis covered larger areas rather than new locations. As such, these denser areas would have a less diffusive impact into non-fibrotic myocardium. This might suggest the hypothesis that fibrosis distributions that are more scattered, as opposed to covering larger areas in the same region, are more consequential for atrial function.

Several clinical studies have performed statistical analysis assessing LA strain metrics, derived from clinical images. Hopman et al. [85] and Chahine et al. [86] both found that reservoir, booster (contractile), and conduit strain were reduced in AF patients compared to healthy controls. In our study, we observed a decrease in booster and reservoir function. However, we did not find the conduit function to change much – except a slight decrease following increased stiffness in fibrotic regions. Our model may not accurately capture changes in conduit function, potentially due to the use of a simplified 0D circulatory model or underestimation of traction force during LV contraction. Another reason might be that we only impose changes in fibrotic regions, while in reality, interstitial fibrosis in AF patients may imply that stiffness is elevated everywhere and not only in fibrotic regions. Fibrosis is not only related to AF; fibrosis levels have been found to be similar for embolic stroke of undetermined source (ESUS) and AF patients [2, 87]. Bashir et al. [88] compared atrial strain metrics for ESUS patients to patients with noncardioembolic stroke. They similarly found that the same metrics (contractile, reservoir, and conduit) were associated with a higher risk of ESUS occurrence, and a higher risk of later detection of AF.

### 4.4 Limitations

Our model representation have several limitations that should be taken into account. Most parameter values were not patient-specific, and those that were still had constraints. In our calibration of patient-specific CV, we assumed healthy baseline simulation values, rather than considering regional fibrotic changes. However, we found CV values to be of low impact among the metrics considered. Thus, this simplification seems reasonable and our main findings should be similar for patient-specific, spatially varying CV values – however, also likely for generic, non-personalized values. Another limitation is that end-diastolic LA volume decreased in the converged solution following ten cardiac cycles, compared to initial volume measurements. The model could be improved by personalizing relevant parameters in the 0D circulatory model to match clinical volume and pressure-related measurements [89, 90]. Other limitations of note include that the model does not incorporate electromechanical feedback [91–94], and the use of a simplified 0D circulatory model to represent hemodynamic feedback, which cannot fully capture fluid-structure interactions [95–97]. In particular, fluid-structure interaction models resolving spatial variations in endocardial pressure might be necessary to resolve secondary interactions involving regional fibrotic changes in myocardial properties.

Our sensitivity analysis only considered a subset of all possible parameter combinations, and we only considered two levels for each parameter. Furthermore, there might other influential parameters parameters not considered in our study that also are altered in cardiac fibrosis. Finally, the results of any sensitivity analysis are always influenced by the intervals selected for each parameter. Our study is limited by the scarcity of experimental data on the relative change in each fibrosis-associated parameter. Despite these limitations, we believe the directionality of each parameter is within reasonable physiological range; future experimental studies might refine exact parameter changes.

A clear limitation of our study is that we only consider three patients, which is very limited especially compared to clinical studies that typically involve hundreds of patients. Data collection involving combined MRI scans and CARTO 3D anatomical mapping for the same patients is logistically challenging, while model simulations involving all combinations is time-consuming. Nevertheless, extending the analysis to a larger cohort would both increase the confidence behind our findings and allow for more granular analysis. Next, our analysis only considered changes in fibrotic regions as determined by LGE, which is calculated and normalized to non-fibrotic myocardium. The approach is not suitable to capture regions of interstitial fibrosis, which is also generally observed among AF patients [5]. As such, there is also remodeling taking place in non-fibrotic regions. This was not taken into account in our analysis, which focused on remodeling in fibrotic regions only.

### 4.5 Future work

This study’s findings warrant future research on quantifying I_CaL_, I_K1_, and myocardial stiffness. Clinical and experimental studies are needed to validate the results presented here, while future computational studies can build on our results. In particular, several clinically drugs used for treatment of atrial fibrillation contains I_K1_-blocking agents [98], while blockers of atrial-specific potassium channels (including Ca^2+^-activated K+ channels of small conductance; SK channels) have been suggested as future therapeutic drugs [99, 100]. These might be used as an alternative to other medications, including but not limited to calcium channel blockers [101]. While primarily assessed in terms of arrhythmogenicity, these drugs may also impact intracellular calcium and force generation. An interesting avenue and extension of our work could be exploring the impact of rhythm control medication on atrial function.

Sensitivity analysis studies of the fibrotic LA could extend in several directions. Incorporation of additional parameters (e.g., other ion channels or parameters relevant to interstitial fibrosis in non-fibrotic tissue) could reveal additional influential mechanisms. Wall deformation predicted by electromechanical simulations could also be integrated with spatially resolved analyses of LA flow and coagulation dynamics to assess how fibrotic remodeling affects downstream outcomes such as thrombosis risk [21, 102–104]. In an extended sensitivity analysis, one could include spatiotemporal analysis of blood flow, including parameters associated to atrial morphology [105], blood rheology [106] (e.g., hematocrit and red blood cell aggregation timescale), coagulatory state, or anticoagulation regime [103]. Another interesting approach could be to extend the sensitivity analysis to ventricular models or whole-heart models, considering the impact of ventricular fibrotic remodeling in comparison or in addition to atrial fibrotic remodeling.

Modeling studies with clinical data can yield highly interesting results. It could be interesting to combine image-based strain analysis and model-based analysis taken for the same patients and controls. Such a study would provide detailed insight into where strain analysis deviates from the model, important for model validation. Another interesting avenue could be to combine our modeling framework with regional analysis of fibrosis distributions [107, 108], relating fibrosis burden and variations in spatial patterns to changes in LA function. Computational modeling could also be used to compare the LA function subject to pulmonary vein ablation, following [109, 110], which could be achieved by treating pulmonary veins as a non-conductive and stiffer material; possibly also combined with post-ablation LGE-MRI images. Findings could be related to clinical data reporting on decreased LA function [67, 79, 111, 112] following pulmonary vein isolation.

## 5 Conclusions

In our study, we used a computational model combined with patient-specific LA geometries to analyze the impact of nine parameters related to fibrotic remodeling. Our sensitivity analysis predicted that impairment of I_CaL_ and I_K1_ were most consequential in terms of changes in LA function, having respectively decreased and improved effect. Future research focusing on these could greatly improve our understanding of fibrotic remodeling and its effects on atrial function. We found that reduction in I_CaL_ and I_K1_ had a diffusive effect also impacting non-fibrotic tissue, and that an increase in fibrosis burden was found to produce a comparable moderate reduction in LA function, which could be related to fibrosis density (scattered versus dense). In the future, modeling frameworks combined with larger cohorts could expand our analysis to better elucidate the relationships between fibrosis burden, spatial patterns, and impairment of LA function. Future modeling efforts could also expand to include spatiotemporal analysis of thrombogenic risk subject to changes induced by fibrotic remodeling, ultimately improving risk assessment and prevention strategies.

## Abbreviations and acronyms

AF: atrial fibrillation
B: baseline
CV: conduction velocity
CV_L_: longitudinal CV
CV_T_: transverse CV
EP: electrophysiology/-ical
EAM: electroanatomical mapping
ESUS: embolic stroke of undetermined source
F: fibrotic
FFD: fractional factorial design(s)
I_CaL_: L-type calcium current
I_K1_: inward rectifier potassium current
I_Na_: fast sodium current
LA: left atrium/-al
LGE: late gadolinium enhancement
µ: myofilament binding rate
OFAT: one-factor-at-a-time
PV: pressure-volume
ST_L_: longitudinal myocardial stiffness
ST_T_: transverse myocardial stiffness
T_a_: active tension scaling factor

## A Supplementary Information: Appendix

### A.1 Calibration of electrical stimulus locations and personalized CV values

For each patient, we obtained LA EAM as recorded during the procedure, from which we used the sinus rhythm maps for calibration of personalized CV values. A CARTO system (J&J MedTech) was used to EAM data during the procedure, and the open-source software OpenEP [1] was subsequently used to extract local activation time maps.

We used the EAM data to identify early activation sites by delineating tissue regions activated within the first 5 ms. For all three patients, this resulted in three disjoint regions. Data for Patient 3 displayed earliest activation for the left superior pulmonary vein; however there was no propagation out of this area and as such this was filtered out by thresholding (see gray areas in Fig A3; bordered by red and yellow). Next, we manually identified the corresponding areas on the volumetric geometries derived from the LGE surfaces. These areas were set as electrical stimulus (pacing) locations in all organ-scale simulations performed.

Next, we estimated personalized CV values for each patient by changing CV values such that the total LA activation time predicted by the EP model simulation matched the value extracted from the CARTO data. We used a constant anisotropy ratio of 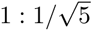 for CV_L_:CV_T_ values, following Zahid et al. [2, 3], such that we only had one varying parameter (CV_L_). We then iteratively ran EP simulations, measured the total LA activation time, and updated the CV values until the total activation time matched the total activation time recorded using EAM data with a less than 1 ms difference.

The algorithm (the iterative process) used to determine optimal CV values for each patient is written out below. In this, the input parameters were a patient-specific geometry, electrical stimulus locations, and total activation time *totalacttime_EAM_* derived from the EAM data as input. Optimized CV values were given as output. That EP simulation corresponded to a full 3D simulation of the electrical propagation for the given geometry over one cardiac beat. The increase and decrease of *CV_L_* was determined manually, for simplicity. While this could also be determined by appropriate incremental increases, we found it easier to do gradual changes manually; with only three cases, this was less work than determining an appropriate scheme for gradual approximation. We did not change any parameters in the fibrotic regions of the patient geometry, assuming that the difference between fibrotic and non-fibrotic configurations would be minor.

#### Algorithm 1 Patient-specific CV calibration

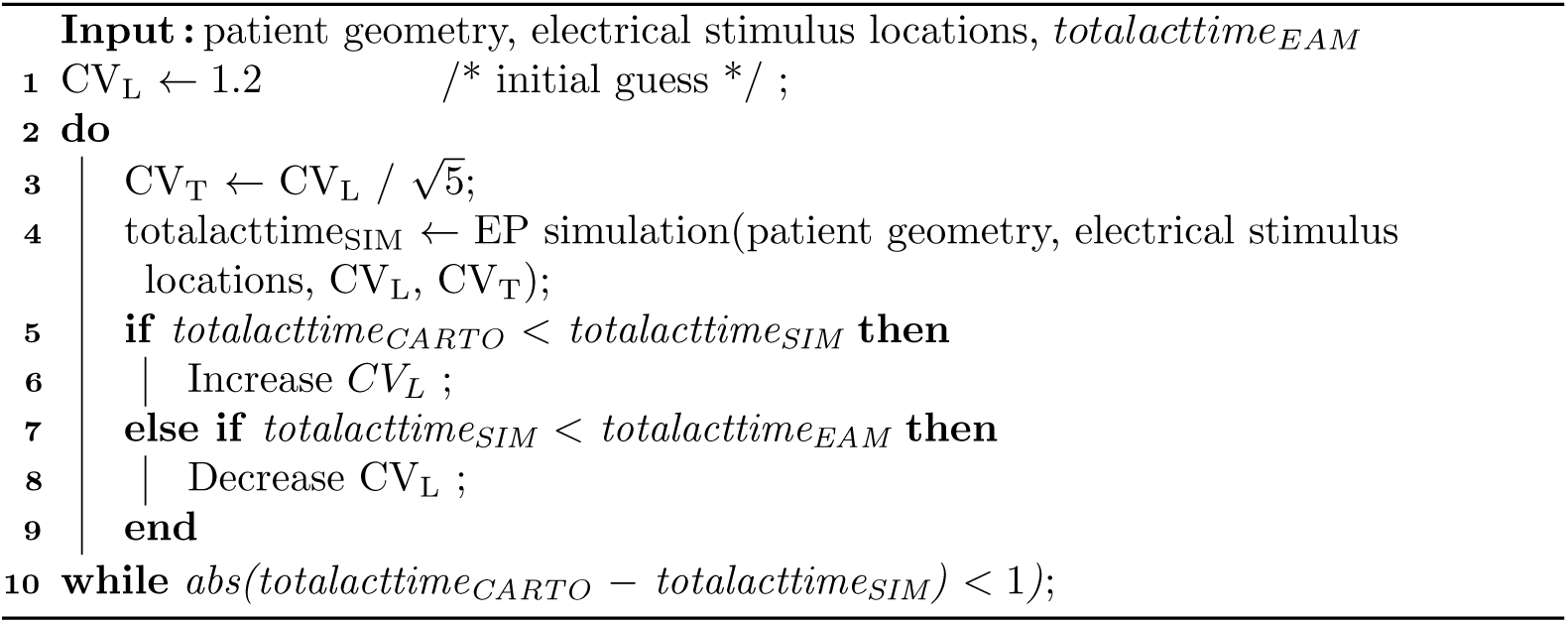

Fig A1–A3 display spatial plots used for determination of earliest activation (pacing) locations, from three different angles. For all three figures, in (A) we display the activation time derived from the EAM (CARTO) data. In (B) we show the same data, but with colors adjusted to highlight the first 5 ms of activation only. In (C) we show the locations we picked on the meshes used for our simulations. The electrical stimulus location was determined manually by matching the area determined by the 5 ms activation area as closely as possible, based on landmarks and geometrical features observed across both geometries. In (D) we display the activation times simulated with our EP model, using electrical stimulus locations from (C) and with optimal CV values as listed in Table 1 in the main manuscript (i.e., healthy baseline values); as well as in Table A1.

### A.2 Patient-specific CV values subject to fibrotic changes

In Table A1, we include CV values imposed from all combinations of reduced and baseline values of all EP parameters considered in our sensitivity analysis schemes. Baseline CV values were calibrated on a patient-specific basis, as previously described. Fibrotic values corresponding to reductions in the longitudinal and transverse directions were calculated by dividing the values by 0.4461 and 0.3526, respectively, as described in the main manuscript. Next, we used open-source software *tuneCV* functionality [4] from the *openCARP* project [5] to estimate the corresponding conductance values for all four values (CV_L_ and CV_T_; non-fibrotic and fibrotic). The conductance values, combined with altered ion channel conduction values, were then used to calculate the corresponding changes in CV values.

**Table A1.** CV values for all combinations of baseline/reduced conductance and baseline/reduced longitudinal/transverse CV for Patient 1 (P_1_), Patient 2 (P_2_), and Patient 3 (P_3_).

|  |  | Baseline CV |  |  | Reduced CV |  |  |
| --- | --- | --- | --- | --- | --- | --- | --- |
| | | $P_1$ | $P_2$ | $P_3$ | $P_1$ | $P_2$ | $P_3$ |
| Conduction values at baseline | $CV_L$ | 1.190 | 1.900 | 0.835 | 0.782 | 1.249 | 0.549 |
| | $CV_T$ | 0.532 | 0.848 | 0.373 | 0.277 | 0.442 | 0.194 |
| $I_{Na} \times 0.6$ | $CV_L$ | 0.947 | 1.532 | 0.653 | 0.610 | 0.995 | 0.418 |
| | $CV_T$ | 0.403 | 0.664 | 0.276 | 0.199 | 0.331 | 0.157 |
| $I_{CaL} \times 0.5$ | $CV_L$ | 1.186 | 1.897 | 0.828 | 0.775 | 1.245 | 0.539 |
| | $CV_T$ | 0.520 | 0.841 | 0.359 | 0.261 | 0.429 | 0.177 |
| $I_{K1} \times 0.5$ | $CV_L$ | 1.221 | 1.930 | 0.868 | 0.815 | 1.280 | 0.583 |
| | $CV_T$ | 0.565 | 0.880 | 0.408 | 0.312 | 0.476 | 0.230 |
| $I_{Na} \times 0.6, I_{CaL} \times 0.5$ | $CV_L$ | 0.942 | 1.529 | 0.645 | 0.601 | 0.990 | 0.405 |
| | $CV_T$ | 0.390 | 0.656 | 0.258 | 0.178 | 0.316 | 0.132 |
| $I_{Na} \times 0.6, I_{K1} \times 0.5$ | $CV_L$ | 0.980 | 1.563 | 0.689 | 0.646 | 1.028 | 0.456 |
| | $CV_T$ | 0.442 | 0.700 | 0.316 | 0.240 | 0.370 | 0.176 |
| $I_{CaL} \times 0.5, I_{K1} \times 0.5$ | $CV_L$ | 1.217 | 1.927 | 0.862 | 0.809 | 1.276 | 0.575 |
| | $CV_T$ | 0.557 | 0.874 | 0.397 | 0.300 | 0.466 | 0.218 |
| $I_{Na} \times 0.6, I_{CaL} \times 0.5, I_{K1} \times 0.5$ | $CV_L$ | 0.975 | 1.560 | 0.682 | 0.638 | 1.023 | 0.446 |
| | $CV_T$ | 0.431 | 0.692 | 0.303 | 0.227 | 0.359 | 0.162 |

### A.3 Average reported metrics for all FFD combinations

In Fig A4 and A5, we include average results for all runs in the FFD in a combined heatmap/tabular format. In this, we first normalized all reported metrics relative to baseline values for each patient individually; then took the average across the three patients.

Notably, in Combinations 9–16 and 25–32 imposed an impairment of I_CaL_. In reported metrics, this is reflected by an overall decrease in A-loop area, booster function, reservoir function, and upstroke pressure difference. Combinations 4–8 and 21–24 involved an impairment of I_K1_ but not of I_CaL_, in which there was an overall increase in metric values. Combinations 13–16 and 29–32 involved an impairment of both, and there was a slight mitigating effect among these compared to Combinations 9–13 and 25–28.

### A.4 Main effect for elevated fibrosis

Main effect plots for all FFD combinations are included in Fig A6, showcasing original versus 50% elevated fibrosis. In both original and elevated fibrosis, we here compared pairwise combinations of all fibrotic factors. Plots corresponding to factor found to be statistically significant are also included in the main paper (Fig. 10).

Comparing original to elevated fibrosis levels, a drop was observed across most metrics (A-loop area, booster function, reservoir function, and upstroke pressure difference), while conduit function did not change much. However the drop was consistent for B and F groups in all but following impairment of I_ICaL_, in which the elevated fibrotic difference was higher (the slope steeper) than in the original fibrosis difference.

---

**Fig A1.**
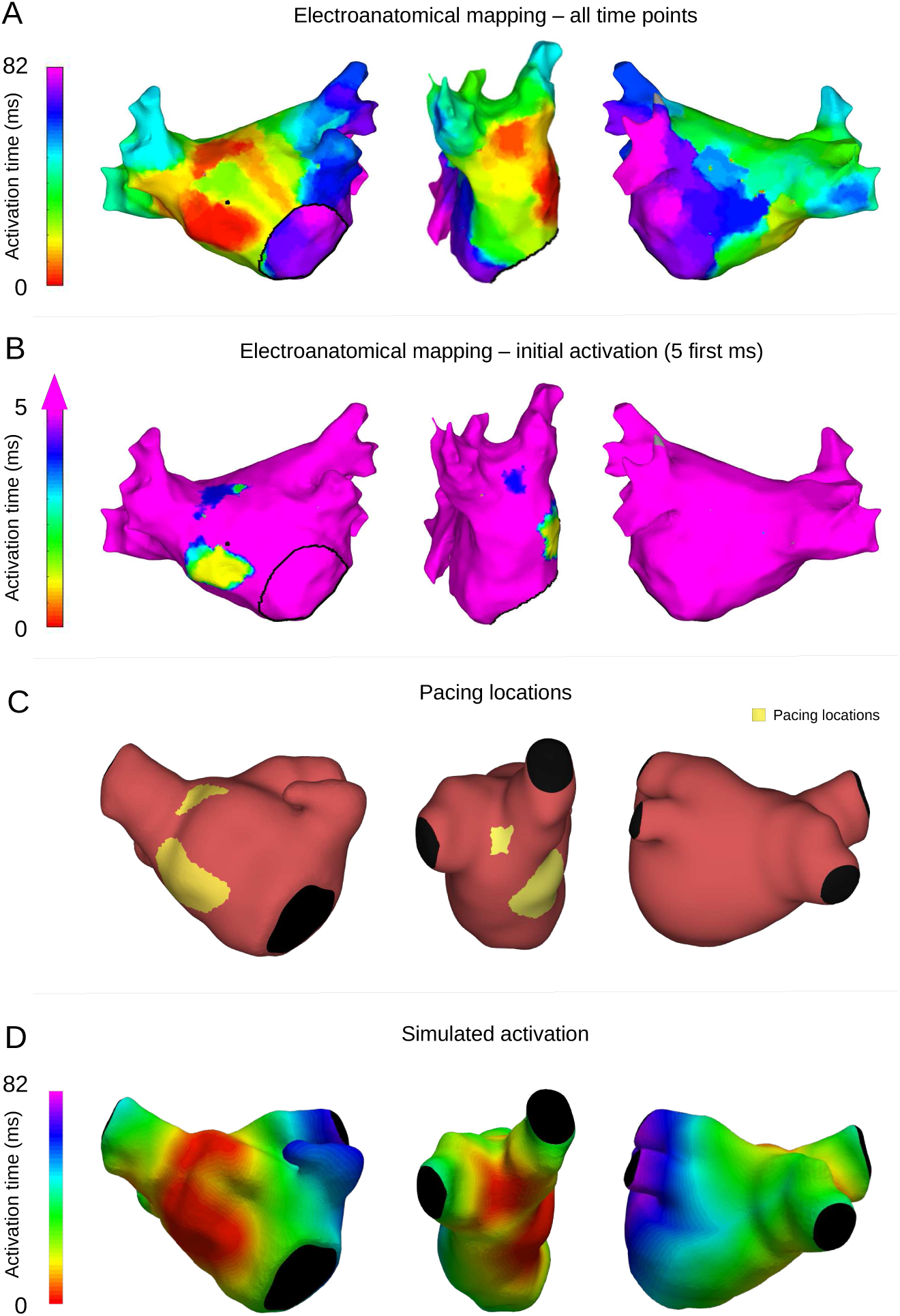
Patient 1 activation times and electrical stimulus locations. (A) Activation times derived from the EAM (CARTO) data, (B) activation times for the first 5 ms, (C) pacing locations, and (D) simulated activation times for Patient 1.

**Fig A2.**
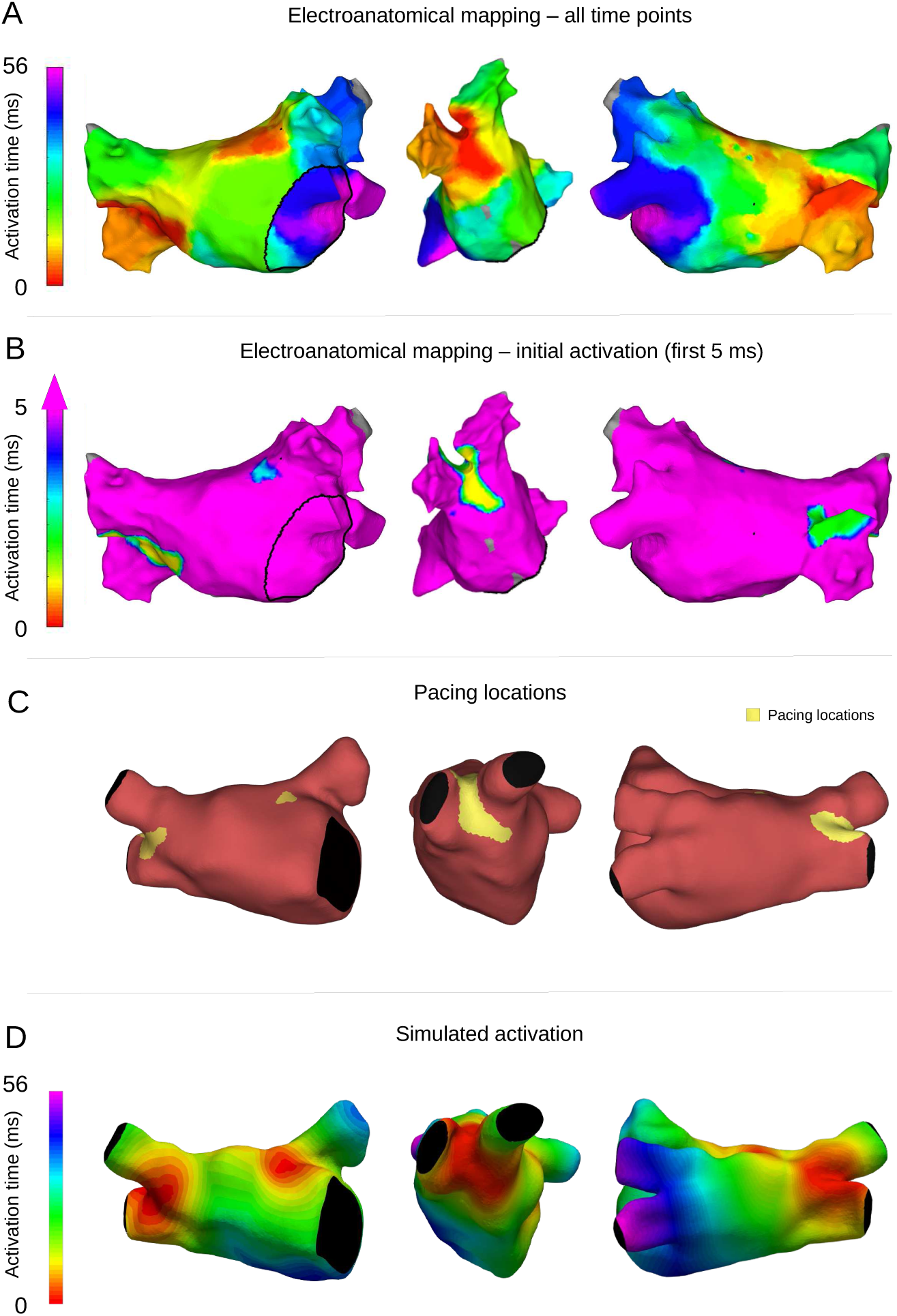
Patient 2 activation times and electrical stimulus locations. (A) Activation times derived from the EAM (CARTO) data, (B) activation times for the first 5 ms, (C) pacing locations, and (D) simulated activation times for Patient 2.

**Fig A3.**
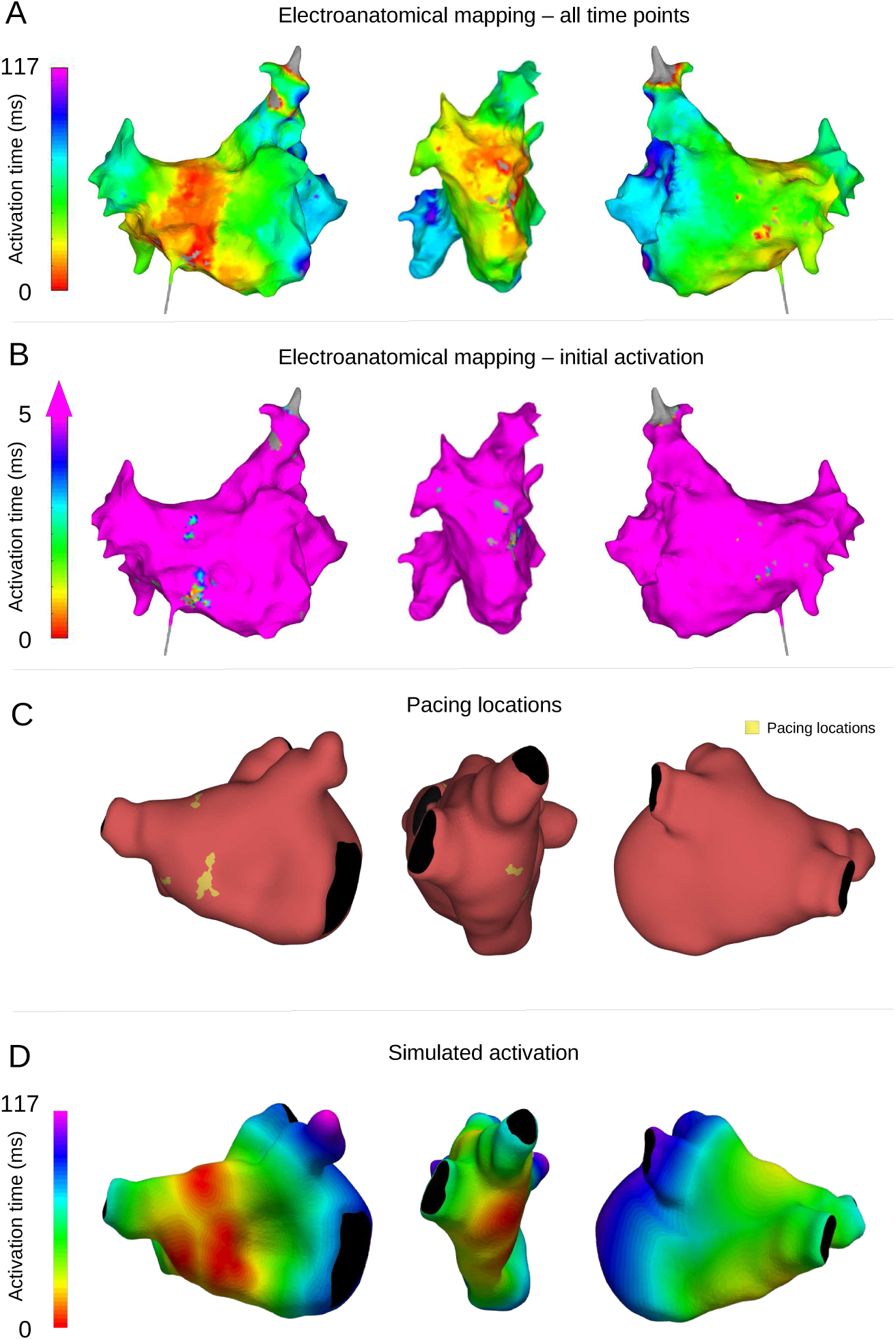
Patient 3 activation times and electrical stimulus locations. (A) Activation times derived from the EAM (CARTO) data, (B) activation times for the first 5 ms, (C) pacing locations, and (D) simulated activation times for Patient 3.

**Fig A4.**
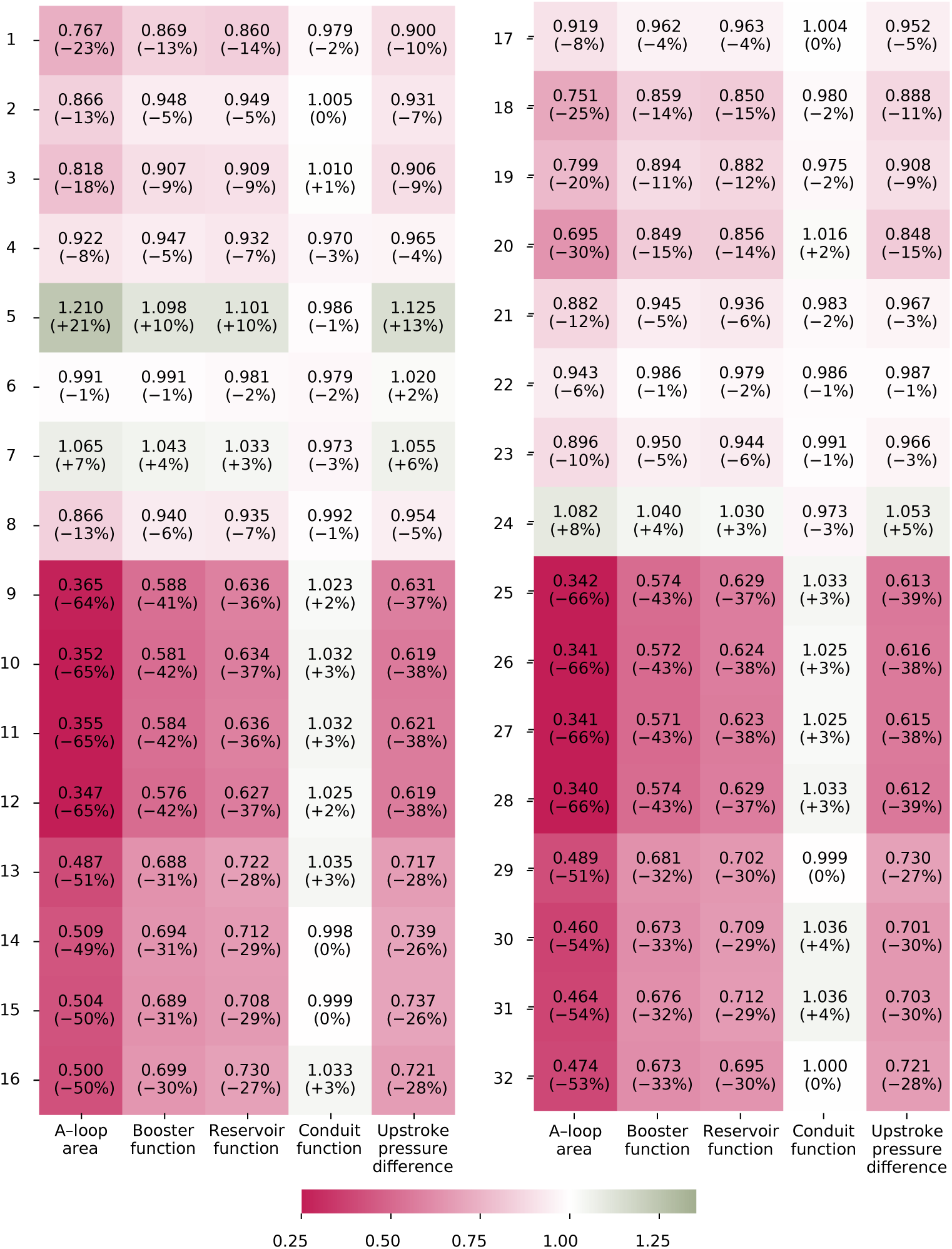
Metrics for FFD Combinations 1–32; original fibrosis burden. Average (across all three patients) normalized (relative to baseline values) metrics for each parameter combination. Corresponding percentage change is listed in parenthesis.

**Fig A5.**
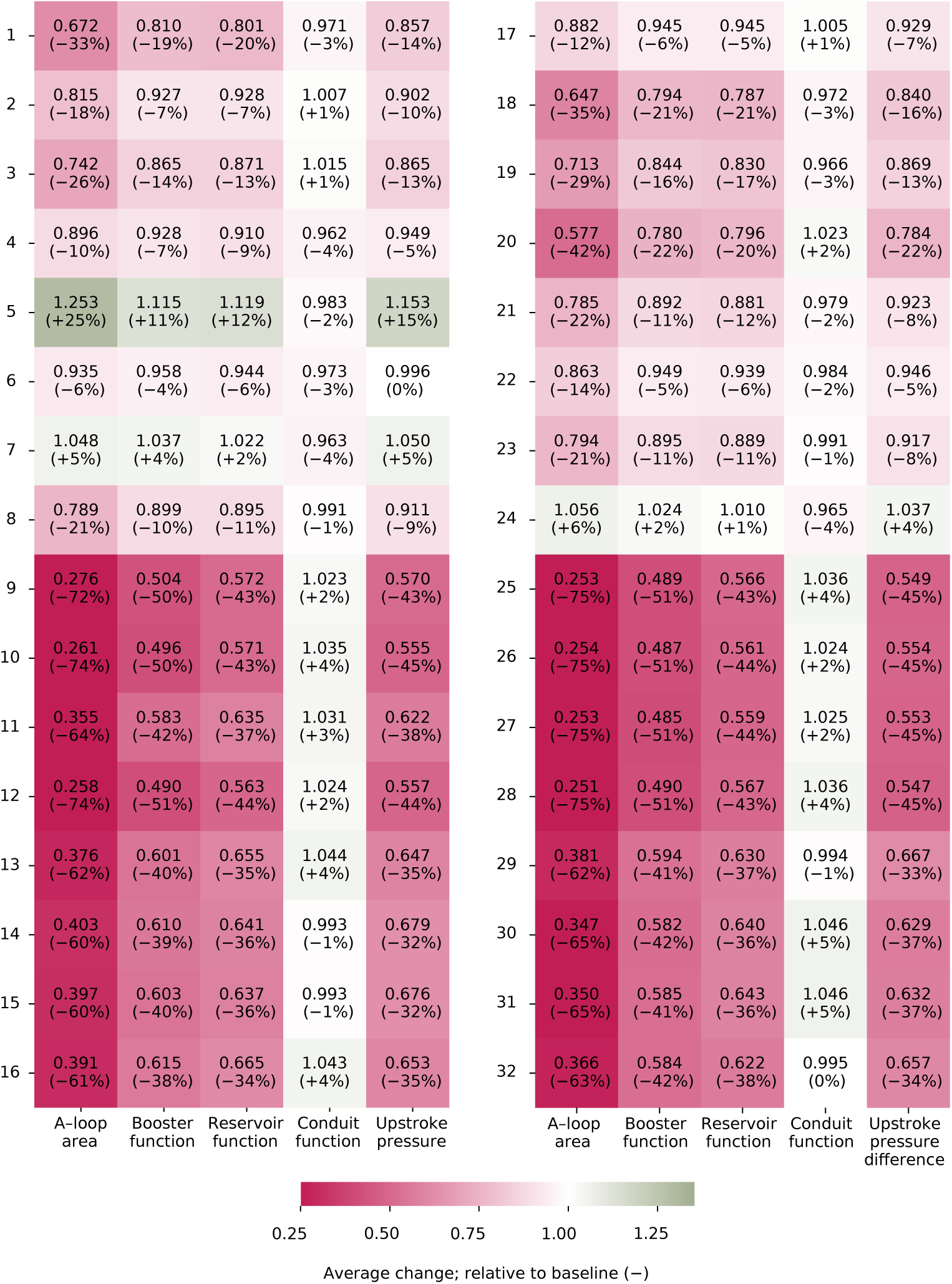
Metrics for FFD Combinations 1–32; 50% synthetically elevated fibrosis burden. Average (across all three patients) normalized (relative to baseline values) metrics for each parameter combination. Corresponding percentage change is listed in parenthesis.

**Fig A6.**
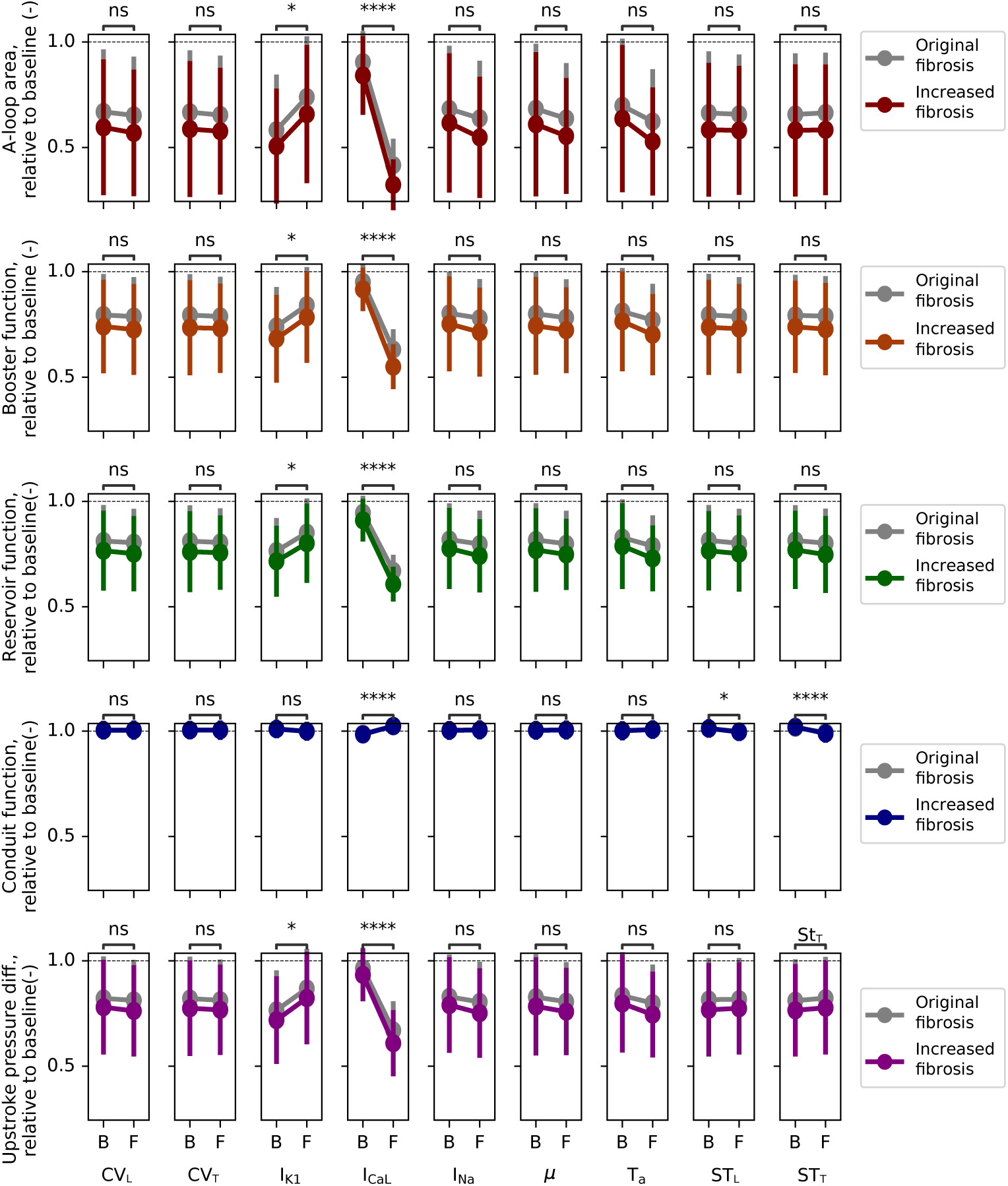
FFD main effect plots; original and 50% synthetically elevated fibrosis. Error bars indicate standard deviation, and gray plots represent original fibrosis levels (same underlying data as in Fig 8 in the main manuscript). Comparisons for significant differences were performed for elevated fibrosis simulations only.

